# Development of a LRRC15-Targeted Radio-Immunotheranostic Approach to Deplete Pro-tumorigenic Mechanisms and Immunotherapy Resistance

**DOI:** 10.1101/2024.01.30.577289

**Authors:** Claire M Storey, Mohamed Altai, Katharina Lückerath, Wahed Zedan, Henan Zhu, Marija Trajkovic-Arsic, Julie Park, Norbert Peekhaus, Jens Siveke, Henrik Lilljebjörn, Diane Abou, Haley Marks, Enna Ulmert, Hans Lilja, Alexander Ridley, Marcella Safi, Constance Yuen, Susanne Geres, Liqun Mao, Michael Cheng, Johannes Czernin, Ken Herrmann, Laurent Bentolila, Xia Yang, Thoas Fioretos, Thomas Graeber, Kjell Sjöström, Robert Damoiseaux, Daniel Thorek, David Ulmert

## Abstract

Leucine-rich repeat containing 15 (LRRC15) has emerged as an attractive biomarker and target for cancer therapy. We have developed a humanized monoclonal antibody (mAb), DUNP19, that specifically binds to a phylogenetically conserved LRRC15 epitope and is internalized by target-expressing cancer and stromal cells. In xenograft mouse models, Lutetium-177 labeled DUNP19 ([^177^Lu]-DUNP19) enables non-invasive imaging and precise radiotherapy to LRRC15-expressing cancer cells and murine cancer-associated fibroblasts (CAFs), halting tumor progression and prolonging survival with minimal toxicity. Transcriptomic analyses of [^177^Lu]-DUNP19-treated tumors reveal a loss of pro-tumorigenic mechanisms, including a transforming growth factor beta (TGFβ)-driven and LRRC15+ signature associated with immunotherapy resistance. Together, these results demonstrate that radio-theranostic targeting of LRRC15 with DUNP19 is a compelling precision medicine platform for image-guided diagnosis, eradication, and reprogramming of LRRC15+ tumor tissue that drives immuno-resistance and aggressive disease.

**SIGNIFICANCE:** We introduce a pioneering LRRC15-guided radio-theranostic approach integrating clinical imaging and radioimmunotherapy. Our strategy utilizes a mAb, DUNP19, to target LRRC15-expressing cancer cells and fibroblasts, demonstrating significant tumor reduction, prolonged survival, and reversal of TGFβ-driven treatment resistance. This approach offers a promising strategy for improving outcomes in aggressive cancers.

## INTRODUCTION

Advancements in the field of immuno-engineering have catalyzed the development of potent antibody-based treatments. However, these therapies are currently applicable towards a narrow subset of malignancies (1). A persistent challenge remains in identifying widely expressed surface antigens for therapy-resistant and metastasized solid tumors (2). The integration of antibodies targeting these biomarkers, alongside advancements in radiochemistry and non-invasive imaging technologies for visualization of radiolabeled antibody distribution, holds substantial promise. This synergy provides a basis for radio-theranostics; selecting patients via imaging that could benefit from treatment and using the same antibody armed with cytotoxic radionuclides for therapy (3). Such an approach could revolutionize cancer treatment, expanding its reach to address a broader spectrum of therapy-resistant and disseminated cancers.

Leucine-Rich Repeat Containing 15 (LRRC15) is a transmembrane protein expressed in TGFβ-driven cancer-associated fibroblasts (CAFs) and cancer cells of mesenchymal stem cell origin, including sarcomas and glioblastomas (4, 5). Although LRRC15 lacks an obvious intracellular signaling domain, recent evidence suggests a role in Wnt/β-catenin signaling pathway activation to promote invasion and metastasis (6, 7). In tumor tissue, the presence of LRRC15+ cells correlates with resistance to immune checkpoint blockade, increased risk for metastasis, and lower survival rates, underscoring LRRC15’s role as an immunomodulator governed by the TGFβ pathway (4, 8-9). Notably, LRRC15 has little or no expression in healthy tissue, making the protein a highly promising target for therapeutic intervention.

Here, we describe the development of a highly specific monoclonal antibody (mAb) targeting LRRC15 (designated as DUNP19) that exhibits high specificity for a phylogenetically conserved epitope present on both human and murine LRRC15, and, upon binding to target cells, rapidly internalizes. We exploit DUNP19’s rapid internalization profile by labeling the mAb with both diagnostic and cytotoxic radionuclides, transforming it into a dual-purpose agent for use in non-invasive imaging and in therapeutic applications. The potential of this technology extends to personalized treatment strategies and dose planning, maximizing the therapeutic index for individual patients.

For diagnostic and therapeutic purposes, we functionalized DUNP19 with positron emitting Copper-64 (diagnostic [^64^Cu]-DUNP19) and beta particle emitting Lutetium-177 (therapeutic [^177^Lu]-DUNP19). As a beta-particle emitter, Lutetium-177 has the potential to cause single-stranded DNA breaks across a span of 10-50 cell diameters. We hypothesized that the crossfire irradiation by [^177^Lu]-DUNP19 could address the challenges presented by the heterogeneous expression of LRRC15 within lesions, especially in larger solid tumors. Our studies, carried out in a range of tumor models with LRRC15+ cancer cells and LRRC15+ CAFs, and in models with LRRC15 expression restricted to CAFs, demonstrate that DUNP19 selectively accumulates in LRRC15+ cells after systemic injection.

Our approach also capitalizes on the deposition of radiation in LRRC15+ cells to eradicate adjacent LRRC15-null tumor tissue and is distinct from classical antibody-drug-conjugates for which target heterogeneity is a limitation. Importantly, systemic treatment with [^177^Lu]-DUNP19 resulted in significant therapeutic efficacy and survival in tumor-bearing mice. In addition, transcriptomic profiling of [^177^Lu]-DUNP19-treated tissues revealed progressive loss of TGFβ-driven genomic signatures associated with malignant disease aggressiveness and immunotherapy resistance. Taken together, these findings highlight the potential of DUNP19 as a radio-theranostic modality for non-invasive detection, targeting, and reprogramming of immuno-suppressive gene signatures within cancer cells and the tumor microenvironment.

## RESULTS

### LRRC15 can be Targeted Utilizing the mAb DUNP19

Our initial investigations tested the specificity of DUNP19 and demonstrated binding to both human and murine recombinant LRRC15 with high affinity (Supp. Fig. 1A). Specific interaction of DUNP19 with LRRC15 was further substantiated by immunoprecipitation of the LRRC15 protein from glioblastoma (U118MG) cell lysates with DUNP19-coated magnetic beads (Supp. Fig. 1B). We next characterized the binding of DUNP19 to LRRC15 across a wide range of cancer cell lines from a range of lineages including melanoma (RPMI7951), glioblastoma (U118MG) and osteosarcoma (HuO9, SAOS2), selecting cell lines based on LRRC15 gene expression data in publicly available databases: the EMBL-EBI expression atlas, Harmonizome 3.0, COSMIC, and the Cancer Cell Line Encyclopedia (CCLE) database (10). Despite selecting cell lines that exhibited high *LRRC15* expression at the RNA level, not all were found to have detectable protein on the cell surface. Among those that did, DUNP19 retained picomolar affinity for cell-surface target antigen (EC50 = 0.83 – 222.22 pM) (Fig. 1A). In addition, a positive correlation was noted between the number of LRRC15 molecules on the cell surface and *LRRC15* mRNA levels (R^2^ = 0.567) (Fig. 1B). Specific binding to LRRC15 on the cell surface was further confirmed by confocal microscopy of AF647-labeled DUNP19 (Fig. 1C). Next, we studied cellular internalization of DUNP19. Antibody internalization enhances the retention time of the delivered radionuclide, thereby increasing both image contrast and absorbed doses of therapeutic radiation (11). Confocal microscopy studies illustrated that DUNP19 is rapidly internalized by LRRC15 expressing cells (Fig. 1D,E). Interestingly, internalization rates for DUNP19 were contingent on the quantity of available molecules; faster kinetics were observed in cells with a higher abundance of LRRC15 molecules (HuO9: 1284.74 ± 63.17 molecules per um^2^, 132.06 ± 10.14 minutes; SAOS2: 684.89 ± 100.67 molecules per um^2^, 145.62 ± 15.18 minutes; Fig. 1D). Conjugation chemistry did not impact DUNP19; the internalization rate of the chelate-conjugated antibody (CHX-A”-DTPA-DUNP19) was analogous to that of the unconjugated antibody (Supp. Fig. 2A). Furthermore, time-resolved cellular assays indicated that ^177^Lu-radiolabeling did not affect affinity (U118MG: Kd = 301 ± 39 pM; HuO9: 117 ± 46 pM; SAOS2: 56 ± 27 pM; RPMI-7951: 25 ± 0.1 pM; Fig. 1F).

**Figure 1.**
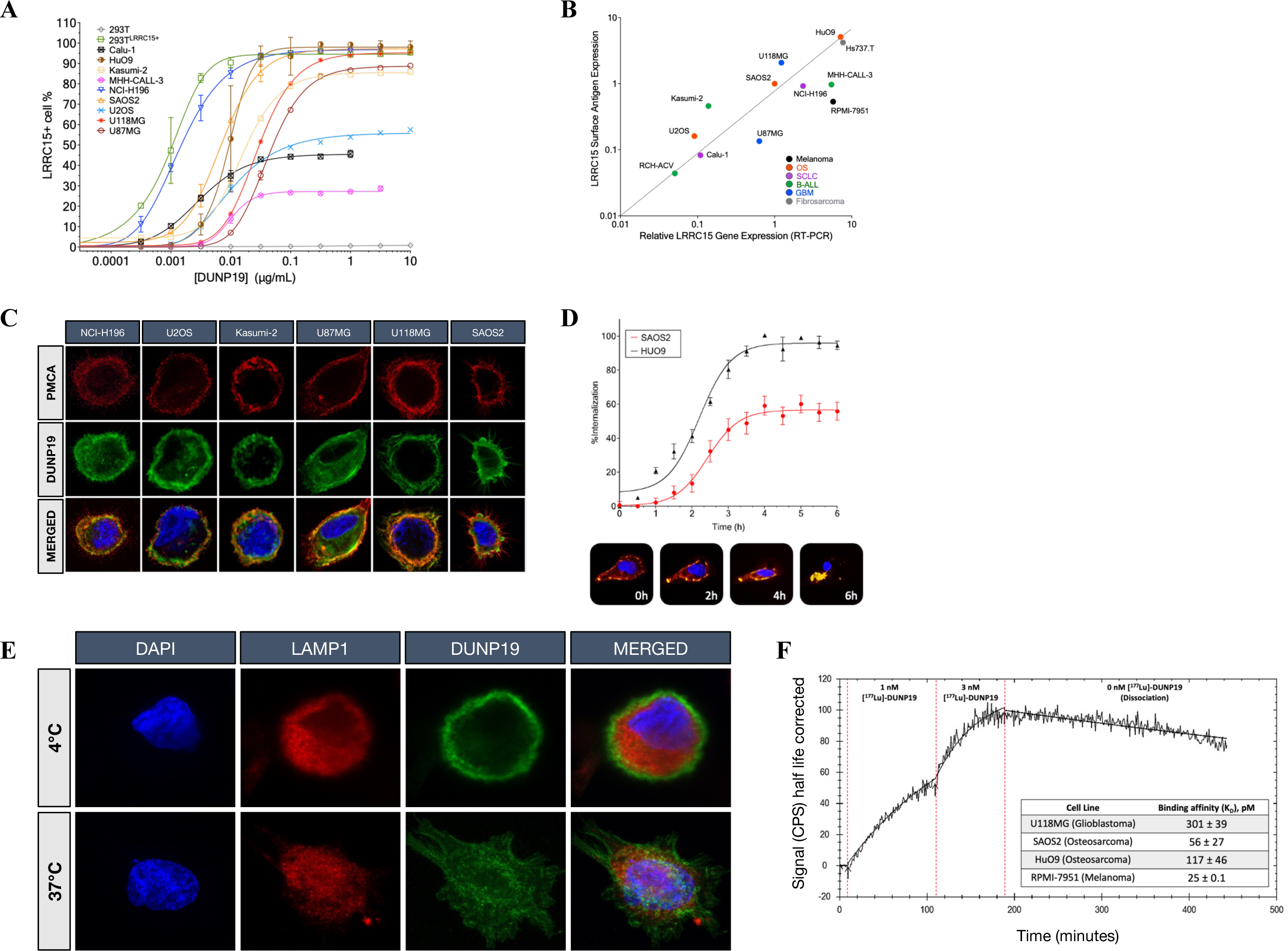
*In vitro* evaluation of DUNP19’s binding profile to LRRC15. **A.** Flow cytometry demonstrates that DUNP19 binds with picomolar affinity to LRRC15 molecules on various cell lines, exhibiting distinct expression levels and tissue origins. Detection of LRRC15 was based on the assessment of antigen-antibody equilibrium. **B.** Analysis across a diverse range of cell lines unveils correlations between *LRRC15* mRNA expression and the abundance of LRRC15 molecules bound by DUNP19. **C.** Confocal microscopy of various cell lines incubated with AlexaFluor647-labeled DUNP19 at room temperature, followed by staining for plasma membrane-associated calcium ATPase (PCMA) and DNA (DAPI). Images reveal that DUNP19 binding corresponds with LRRC15 expression levels and co-localizes with LRRC15. **D.** Internalization rates of AlexaFluor647-labeled DUNP19 at 37°C were examined in LRRC15-expressing HuO9 and SAOS2 cells using live confocal microscopy. The endocytic process of DUNP19 is accelerated in cells with higher LRRC15 abundance. **E.** Confocal microscopy of SAOS2 cells incubated with AlexaFluor647-labeled DUNP19 at 4°C or 37°C, co-stained for lysosomes (LAMP1) and DNA (DAPI). DUNP19 is exclusively found in the plasma membrane at the lower temperature demonstrating that the rapid endocytosis after binding to LRRC15 is an active, energy-requiring process. **F.** LigandTracer sensorgram of [^177^Lu]-DUNP19 binding to LRRC15-expressing HuO9 cells, measured at 1 nM and 3 nM concentrations. Cell-bound activity, presented as CPS, was used to determine association, dissociation rates, and equilibrium dissociation constants (K_D_). The table shows K_D_ values for various LRRC15-expressing cell lines.

### DUNP19 *In Vivo* PET Can Monitor Tumor-Associated LRRC15 Expression

To evaluate *in vivo* kinetics of DUNP19 in healthy organs and LRRC15-expressing tumors, sequential PET images were acquired of subcutaneous (s.c.) osteosarcoma (SAOS2) bearing mice after intravenous (i.v.) administration of a ^64^Cu-labeled version of the mAb ([^64^Cu]-DUNP19). Tumor accumulation was compared to the clinical bone scanning agent [^18^F]-NaF. Rapid [^64^Cu]-DUNP19 uptake *in vivo* recapitulated *in vitro* internalization, selectivity, and retention in malignant tissue. In contrast, [^18^F]-NaF exhibited limited accumulation in osteogenic tumors, with the bladder showing the highest activity due to urinary excretion (Fig. 2A). These findings suggest that DUNP19-PET can be utilized to noninvasively determine LRRC15 expression and select patients for treatment, improving upon existing FDA-approved PET tracers for bone cancer lesions.

**Figure 2.**
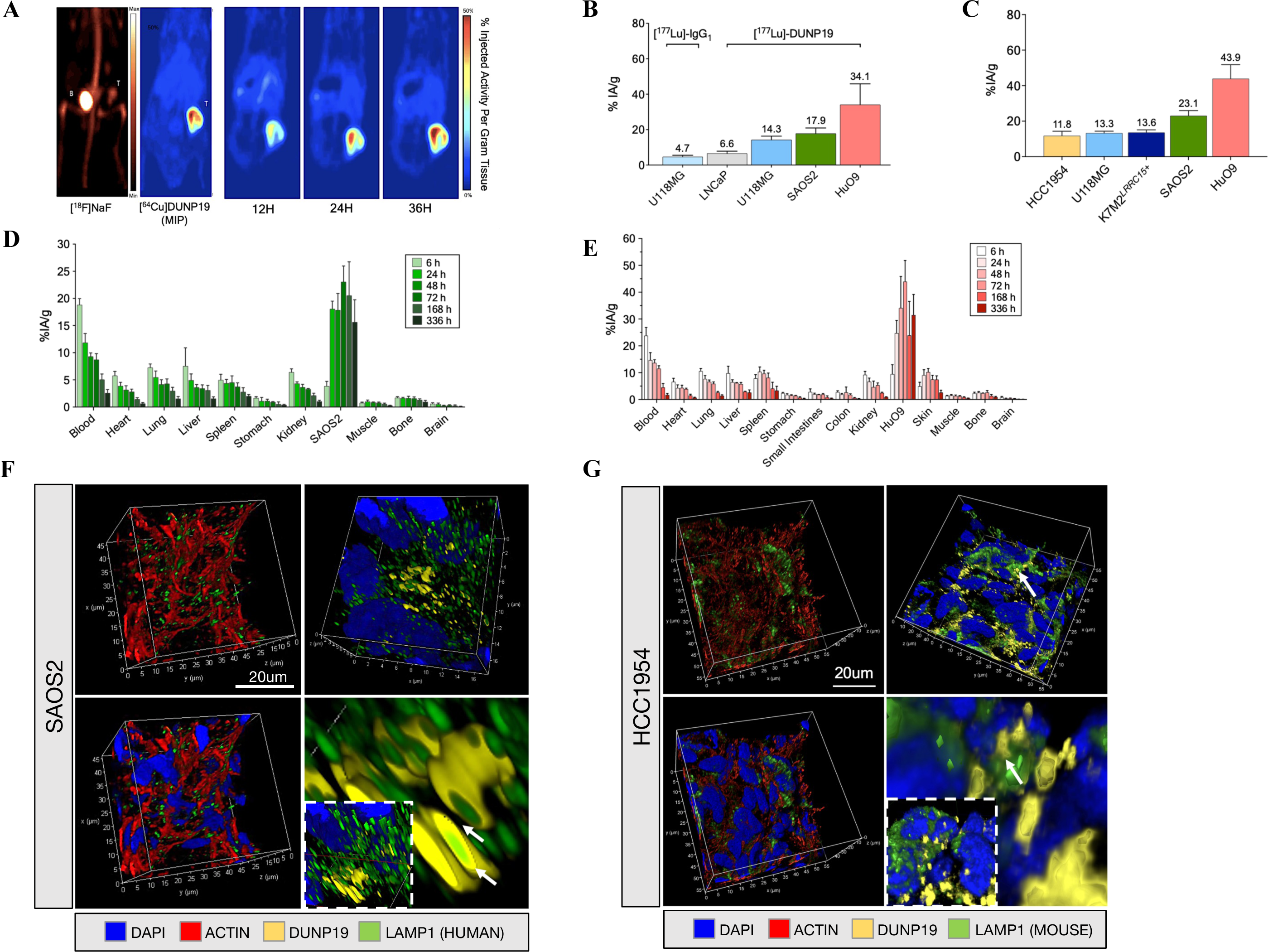
Specificity and biodistribution of DUNP19 in multiple mouse models with LRRC15 expressing tumors. **A.** Representative PET images of s.c. SAOS2 osteosarcoma xenografts obtained at different time points post i.v. administration of [^64^Cu]-DUNP19, highlighting significant tumor-specific uptake with minimal accumulation in normal tissues. In contrast, PET with the clinical bone scanning agent [^18^F]-NaF showed low activity in tumor tissue, with the majority of the tracer dose observed in bone (Bn) and bladder (Bl). **B.** *In vivo* assessment of LRRC15 targeting specificity by [^177^Lu]-DUNP19. At 48 h post i.v. injection, [^177^Lu]-DUNP19 displayed significantly higher uptake (p < 0.001) in LRRC15+ U118MG (blue bar) and HuO9 (red bar) tumors compared to LRRC15-LNCaP tumors (light gray bar). The accumulation of non-specific [^177^Lu]-IgG1 in LRRC15+ U118MG tumors (dark grey bar) was significantly lower than that of [^177^Lu]-DUNP19. **C.** [^177^Lu]-DUNP19 tumor uptake in multiple s.c. tumor models at 72 h p.i., correlating with the LRRC15 expression level in the respective model. **D. + E**. Kinetics of [^177^Lu]-DUNP19 in healthy organs and LRRC15+ SAOS2 and HuO9 osteosarcoma lesions. *Ex vivo* tissue biodistributions of [^177^Lu]-DUNP19 obtained at multiple time points after i.v. injection showed a continuous decline in activity levels in healthy organs, but sustained uptake by malignant lesions. **F. + G.** Microanatomy of tumor tissues obtained from animals treated with fluorescently labeled DUNP19. Confocal images of s.c. SAOS2 (LRRC15+ cancer cells / LRRC15+ CAF) and HCC1954 (LRRC15-cancer cells / LRRC15+ CAF) tumors harvested at 72 h post-i.v. injection of AF594-DUNP19 (yellow). Tumor sections were co-stained for Actin (red), DNA (DAPI, blue) and LAMP1 (lysosomal marker, green). Images show that DUNP19 accumulates in the cellular cytoplasm and co-localized with LAMP1 indicating intracellular trafficking of the mAb to the lysosomal compartments (arrow) after binding membranous LRRC15.

### [^177^Lu]-DUNP19 Exhibits a Favorable Biodistribution

^177^Lu is a clinically relevant beta particle emitter with a half-life of 6.7 days and maximum tissue penetration range of 1.5 mm. Because of the radionuclide’s long path length, ^177^Lu can be used to deliver ionizing radiation to target cells and to target-null cells in close proximity via a crossfire effect (12). Given these characteristics, we hypothesized that [^177^Lu]-DUNP19 could overcome the heterogeneous expression of LRRC15 observed across tumor tissue and stroma (5). First, we evaluated the influence of antibody carrier mass on the biodistribution of [^177^Lu]-DUNP19 in s.c. osteosarcoma (HuO9) tumors. We found that an injected antibody mass of 15-30 µg yielded the optimal tumor-to-normal tissue radioactivity uptake ratios. Next, we systematically examined the biodistribution and pharmacokinetic profile of [^177^Lu]-DUNP19 in a variety of s.c. tumor models originating from diverse malignant tissues. These studies evaluated multiple cancer lineages with various levels of LRRC15 expression and with target expression in distinct tumoral compartments. Tumor models with LRRC15+ cancer cells and CAFs included K7M2*^LRRC15+^*, HuO9 and SAOS2 osteosarcomas (OS) and U118MG glioblastoma (GBM). We also assessed [^177^Lu]-DUNP19 uptake in the HCC1954 breast cancer model, characterized by high LRRC15 expression in CAFs and LRRC15-null cancer cells. (Fig. 2, Supp. Fig. 3, 4).

The accumulation of [^177^Lu]-DUNP19 in tumors peaked at 72 h post-injection (p.i.) (HCC1954: 12.5 ± 2.8 %IA/g [percent injected activity per gram tissue], U118MG: 13.3 ± 1.1 %IA/g, K7M2*^LRRC15+^*: 13.6 ± 1.5 %IA/g, SAOS2: 23.1 ± 2.9 %IA/g, HuO9: 43.9 ± 7.9 %IA/g) and remained consistently elevated at all studied time points throughout the time course, out to 336 h p.i. (Fig. 2B). Retention of [^177^Lu]-DUNP19 steadily decreased in blood and healthy organs after injection, and [^177^Lu]-DUNP19 in blood reflected the expected half-life of a human IgG_1_ in mice, indicating interaction with the murine neonatal fragment crystallizable (Fc) region receptor (FcRn) (13, 14). Retention of [^177^Lu]-DUNP19 in the liver was representative of typical blood volume and metabolic elimination of antibodies. Taken together, these data indicate a favorable biodistribution profile of [^177^Lu]-DUNP19 (Fig. 2D,E).

LRRC15 targeting specificity *in vivo* was further addressed in the s.c. LRRC15+ glioblastoma tumor model, U118MG. Uptake of [^177^Lu]-DUNP19 was compared to nonspecific [^177^Lu]-huIgG_1_, a human IgG_1_ with non-binding complementary-determining regions (CDRs) that had been radiolabeled with ^177^Lu. At 48 h post-i.v. administration, tumor uptake of [^177^Lu]-DUNP19 was significantly higher than [^177^Lu]-huIgG_1_ (14.31 ± 2.01 vs. 4.72 ± 0.82 %IA/g, respectively). Additionally, in LRRC15-s.c. LNCaP tumors, which lack relevant amounts of murine LRRC15 expressing stroma but are highly vascularized, systemic injection of [^177^Lu]-DUNP19 resulted in tumor retention of 6.57 ± 1.28 %IA/g at 48 h p.i. (Fig. 2B). These findings indicate that DUNP19 specifically targets LRRC15-expressing tumor tissue with minimal off-target retention *in vivo*. Accumulation of [^177^Lu]-DUNP19 in LRRC15-LNCaP tumors is likely due to the enhanced permeability and retention effect. This pathophysiological mechanism involves the entrapment of macromolecules >45 kDa by the tumor vasculature, exhibiting a pronounced non-specific uptake of compounds in small animal xenograft tumor models as opposed to malignant lesions observed in humans (14, 15).

Next, we investigated the subcellular localization of DUNP19 in human osteosarcoma (SAOS2 and HuO9), and breast cancer (HCC1954) tumor models following systemic administration. Sections from s.c. tumors collected 72 h after i.v. injection of AlexaFluor647-labeled DUNP19 were co-stained for DNA, actin, and lysosomes (LAMP1) and analyzed by confocal microscopy (Fig. 2F,G, Supp. Fig. 4). Consistent with our *in vitro* findings, DUNP19 co-localized with murine LAMP1 in HCC1954 tumors, and with human LAMP1 in HuO9 and SAOS2 tumors. This co-localization suggests cellular internalization of the antibody after binding with LRRC15 on the plasma membrane of both cancer and stromal cells.

### [^177^Lu]-DUNP19 Radioimmunotherapy (RIT) of Aggressive Osteosarcoma

We then investigated the impact of [^177^Lu]-DUNP19 radioimmunotherapy (RIT) on tumor volume and overall survival in mice bearing HuO9 osteosarcoma tumors. Following a single systemic administration of 30 MBq [^177^Lu]-DUNP19, tumor growth was significantly inhibited (P<0.005, Fig 3A_I_). All mice treated with a single administration of 30 MBq [^177^Lu]-DUNP19 survived through the entirety of the observation period (126 days) for initial tumor volumes in the 150-200 mm³ range. Infiltrative and bulky osteosarcoma have poorer prognoses and reduced treatment options. To test the impact of a larger tumor burden, a second group of animals with greater tumor volume (500-600 mm³) were treated with a single administration of 30 MBq [^177^Lu]-DUNP19. Disease progression was again delayed relative to control, and overall survival was also improved to 115 days (67-126 days). All untreated animals succumbed by day 65 days (range: 58-69 days) (Fig 3A_II_). Thus, irrespective of initial tumor volume, treatment with [^177^Lu]-DUNP19 RIT was effective at significantly reducing disease progression and improving overall survival.

**Figure 3.**
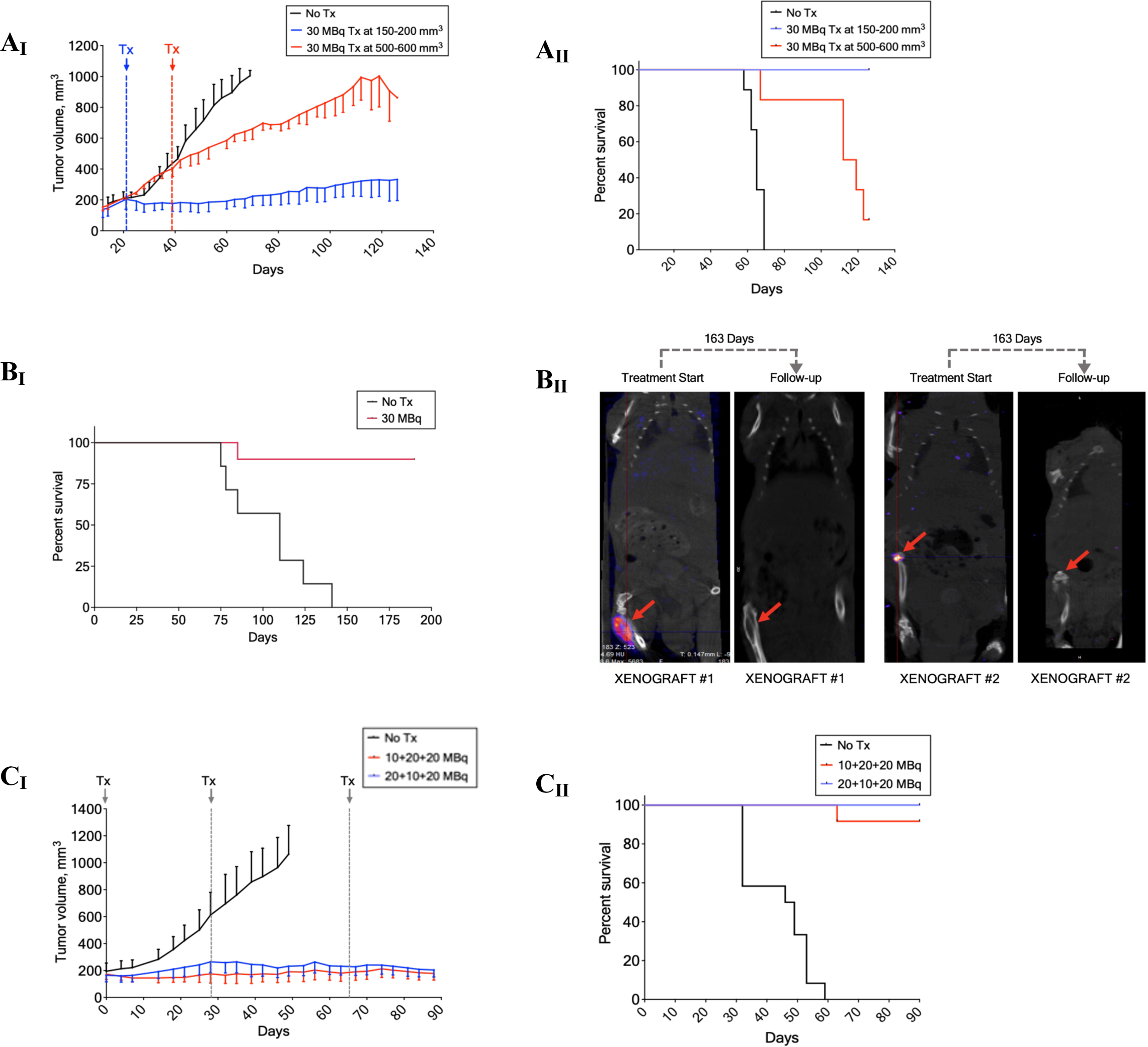
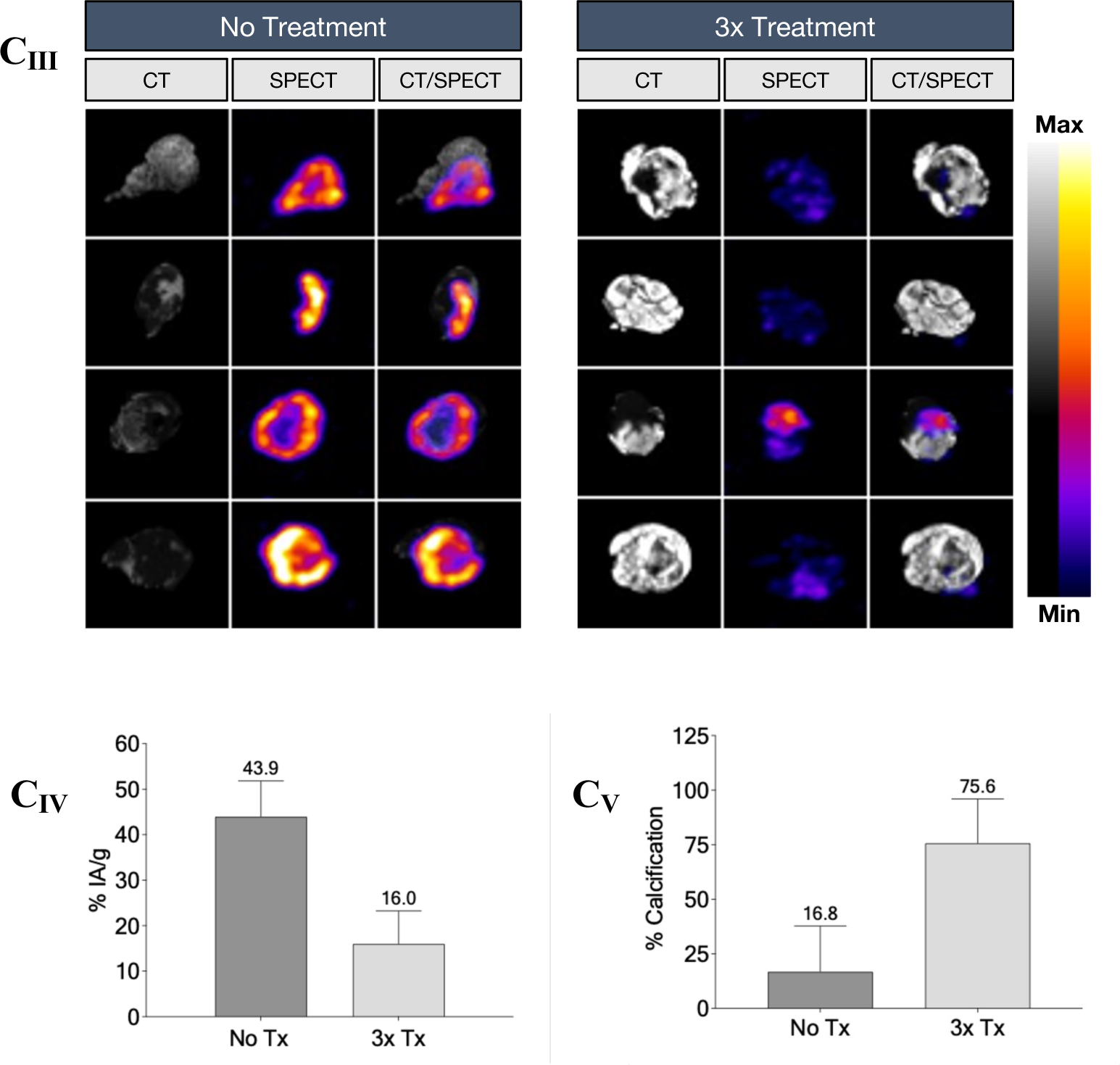
Evaluation of [^177^Lu]-DUNP19 therapy and monitoring in HuO9 osteosarcoma tumors. **A_I-II._** Tumor volumes in mice bearing s.c. HuO9 xenografts (n=10 per arm) were randomized for a single i.v. administration of 30 MBq [^177^Lu]-DUNP19 when tumors reached 203±64 mm3 (blue line; day 21) or 504±152 mm3 (red line; day 39) or received no treatment (black line). The results demonstrated a significant delay in disease progression, with treatment efficacy being influenced by tumor volume (**A_I_**). Kaplan-Meier survival analysis revealed that [^177^Lu]-DUNP19 extended survival, with the impact varying based on the timing of intervention (**A_II_**). **B_I-II_.** Representative coronal SPECT/CT images showing orthotopic HuO9 osteosarcoma tumors (indicated by arrow) after initial (left) and follow-up (right) i.v. administrations of 20 MBq [^177^Lu]-DUNP19. In all treated mice (right) (n=10), no tumor associated uptake was observed at 163 days after treatment **(B_I_)**. Kaplan-Meier survival analysis revealed a significant increase in survival for the [^177^Lu]-DUNP19 treated group during the observed 190-days period (**B_II_**). **C_I-V_**. Mice with s.c. HuO9 osteosarcoma xenografts (n=12 per arm) were randomized for three treatment cycles (red: 10+20+10 MBq; blue: 20+10+20 MBq) of i.v. [^177^Lu]-DUNP19 (at day 0, 32, and 75), resulting in a total administered activity of 50 MBq, or no treatment (black; n=12). Assessment of tumor volumes demonstrated that repeated cycles of LRRC15-RIT effectively inhibit tumor growth (**C_I_**). Kaplan-Meier survival analysis confirmed significantly improved survival for mice randomized for [^177^Lu]-DUNP19 over no treatment (**C_II_**). Four tumors from the treatment and control (non-treatment) arm, harvested 72 hours after administration of an imaging dose of [^177^Lu]-DUNP19 (3 MBq), were imaged ex vivo by SPECT and CT (**C_III_**). Tissue activity levels (%IA/g), assessed by gamma counter and normalized to tissue weight, revealed significantly lower uptake of the antibody in treated vs. non-treated tumors (p < 0.001), reflecting reduction in total LRRC15-expressing cells post-treatment (**C_IV_**). Quantification of radiopacity in CT images showed significantly higher ossification levels in treated vs. non-treated tumor tissues (p < 0.001) (**C_V_**). Together, these findings illustrate that repeated cycles of [^177^Lu]-DUNP19 effectively reduce tissue viability and calcification.

To investigate the efficacy of [^177^Lu]-DUNP19 dosing regimens on tumor growth and survival, we conducted therapeutic fractionated dosing studies. From a translational perspective, this approach is commonly utilized in clinical settings to optimize maximum tolerated dose while reducing dose-limiting toxicities (16-18). Fractionation is also recommended to compensate for the anticipated heterogeneity in RIT dose distribution, particularly in large poorly vascularized tumors with regions of hypoxia (19). Rather than adhering to a predetermined activity and treatment schedule, additional therapeutic doses were given based on recovery from bone marrow toxicity determined by measuring differential blood counts, regrowth of tumor volume, and their effect on animal overall weight (Supp. Fig. 5,6). Mice bearing subcutaneous HuO9 tumors (150-200 mm³) were administered a cumulative activity of 50 MBq in three fractions over a span of 88 days (Fig. 3C_I_). Continuous progression-free survival was sustained in treated animals throughout the study duration, contrasting to untreated animals which exhibited a median survival of 47 days (Fig. 3C_II_).

We further studied LRRC15-targeted radio-theranostics in a translationally relevant orthotopic osteosarcoma model (Fig. 3B-D). HuO9 cells were injected into the left tibia of Balb/c mice and half of the subjects were randomly selected for systemic injection with 30 MBq of [^177^Lu]-DUNP19 23 days after inoculation. Imaging by SPECT at 72 h post-[^177^Lu]-DUNP19 administration revealed LRRC15-specific accumulation of activity at the tumor site (Fig. 3B). Of treated animals, 9 of 10 survived to study endpoint (190 d), while all untreated animals succumbed due to disease-related endpoints (Fig. 3B_I_). Follow-up imaging with [^177^Lu]-DUNP19 was acquired at 163 days after treatment and revealed no accumulation of the theranostic agent at the site of HuO9 tumor inoculation or at other anatomical locations (Fig. 3B_II_). Based on these results, we concluded that the previously detected LRRC15+ tissue had been eradicated.

Radiation is widely known to induce calcifications in sarcomatous processes (20). To investigate this phenomenon in animals bearing HuO9 lesions, we quantified tumor radiopacity and uptake of tumor-associated [^177^Lu]-DUNP19 using SPECT/CT and gamma-spectrometry. Animals treated with [^177^Lu]-DUNP19 and untreated mice received an imaging dose of 3.5 MBq [^177^Lu]-DUNP19 and tumors were harvested 72 hours after injection. Treated tumors exhibited significantly lower tumor-associated [^177^Lu]-DUNP19 activity, coupled with higher levels of tumor calcification (Fig. 3C_III__-V_).

### [^177^Lu]-DUNP19 Therapy is Effective Across a Range of Tumors with Varying LRRC15 Expression Patterns

Given the high expression of LRRC15 on HuO9 osteosarcoma cells (Fig. 1), we also sought to understand how the effects of [^177^Lu]-DUNP19 therapy would change in a tumor of different tissue origin and with lower LRRC15 expression. Mice bearing s.c. tumors from glioblastoma U118MG cells were treated with a cumulative activity of 20 MBq or 30 MBq [^177^Lu]-DUNP19 in two fractions. [^177^Lu]-DUNP19 treatment significantly extended survival (p<0.0001), and therapy prevented further tumor growth as tumor volumes reached a plateau phase around 100-200 mm^3^. All untreated mice succumbed due to disease-related endpoints by 78 days, whereas 11/12 mice (91.67 %) and 12/12 mice (100%) were alive at the end of the observation period in the 20 MBq and 30 MBq treatment groups, respectively (Fig. 4 A,B).

**Figure 4.**
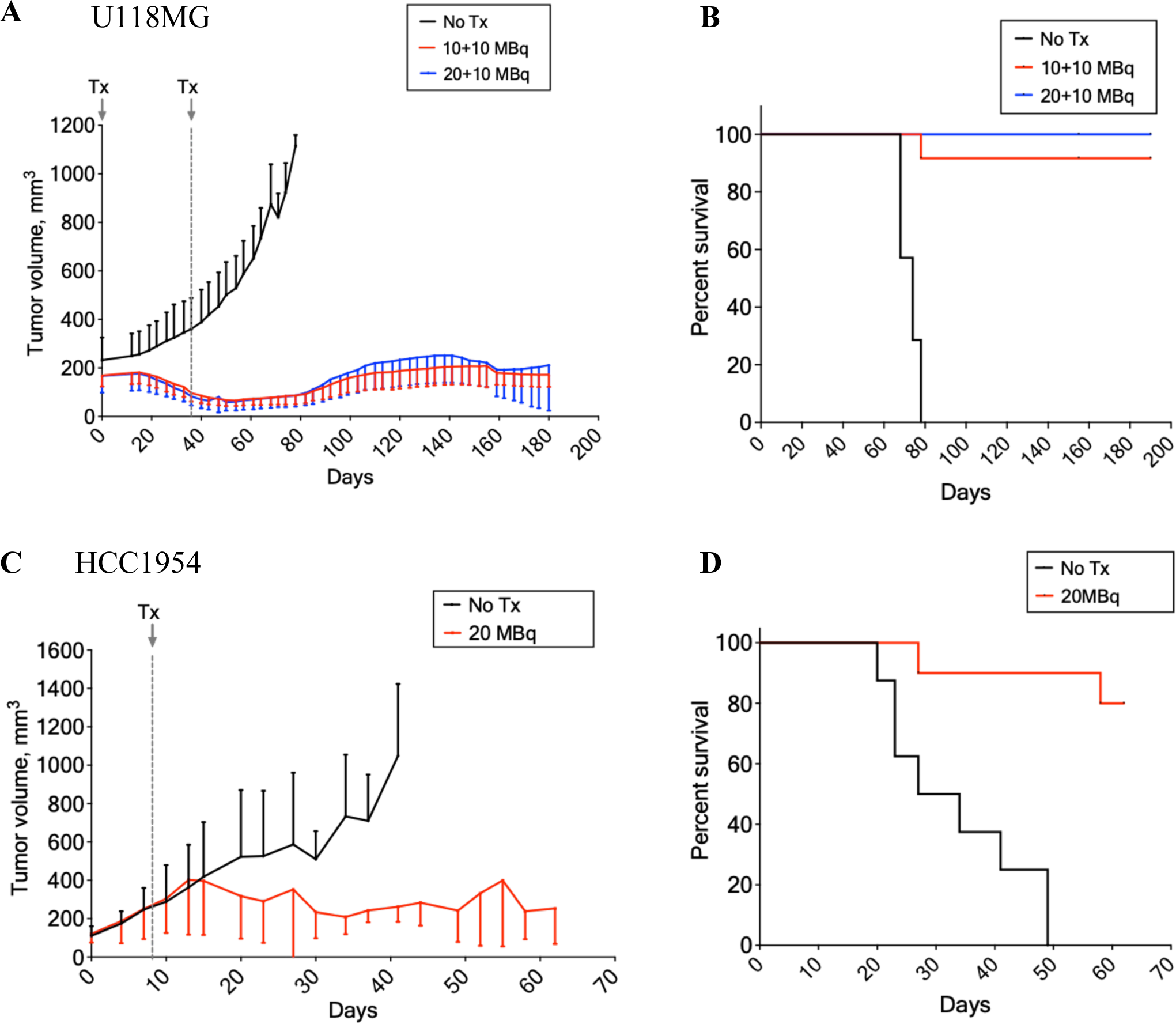
Therapeutic efficacy of [^177^Lu]-DUNP19 in LRRC15-expressing human xenograft models. **A, B.** [^177^Lu]-DUNP19 demonstrates antitumor activity in other cancer indications. BALB/c nude mice bearing s.c. U118MG glioblastoma xenografts (were treated with two fractions of [^177^Lu]-DUNP19 at days 0 and 34 for a cumulative activity of 20 MBq (10+10 MBq, red, n=12) or 30 MBq (20+10 MBq, blue, n=11). Despite lower LRRC15 expression by U118MG tumors, treatment with [^177^Lu]-DUNP19 significantly controlled tumor growth and prolonged survival in both [^177^Lu]-DUNP19 doses (median survival; untreated = 74 days, 20 MBq = not reached, 30 MBq = not reached, p < 0.001). **C, D.** [^177^Lu]-DUNP19 is effective in HCC1954 breast cancer models (LRRC15-cancer cells, LRRC15+ stroma). Results demonstrate delayed s.c. HCC1954 growth in female mice intravenously administered a single dose of [^177^Lu]-DUNP19 (20 MBq; day 7, n=10). **D.** Median survival was not reached for the treated group by the end of the observation period (day 62), while median survival of treated mice was 30.5 days (p < 0.005).

Finally, we evaluated the therapeutic efficacy of [^177^Lu]-DUNP19 in aggressive breast cancer tumors comprising LRRC15-HCC1954 cancer cells and LRRC15+ murine CAFs. Despite LRRC15 expression in stroma only, a single systemic injection of 20 MBq [^177^Lu]-DUNP19 effectively suppressed HCC1954 tumor growth and significantly prolonged median survival compared to untreated mice, where median survival was 30.5 days (p=0.0005; median survival not reached in treated animals). Notably, 80% of the treated mice (8 out of 10) survived until the end of the observation period (Fig. 4C,D). Given the lack of target expressing cancer cells, the response of HCC1954 tumors to [^177^Lu]-DUNP19 is reliant on targeting of LRRC15+ murine CAFs. We presume that the therapeutic impact results from the synergy of two components: the direct induction of DNA damage by beta-particles, affecting both stromal and cancer cells, as well as the ablation of tumor-supporting stromal tissue on which cancer cells depend for growth and survival.

Throughout all studies, treatment was well-tolerated as indicated by stable body weights (Supp. Fig. 5). Administration of [^177^Lu]-DUNP19 resulted in a transient bone marrow suppression, which recovered to baseline levels within 21 days (Supp. Fig. 6).

### LRRC15-targeted RIT Depletes TGFβ-Driven Signature in Tumors

Having demonstrated the significant therapeutic potential of our radio-immunotheranostic platform, we sought to understand the molecular effects of LRRC15-targeted RIT on cancer cells and the tumor microenvironment. Transcripts of bulk RNA-sequencing of tumors harvested at 90-155 days after [^177^Lu]-DUNP19 treatment were aligned to human and murine genomes to identify the transcriptomic profiles of human cancer cells and murine stromal cells. Ambiguous reads were subsequently removed (Fig. 5A).

**Figure 5.**
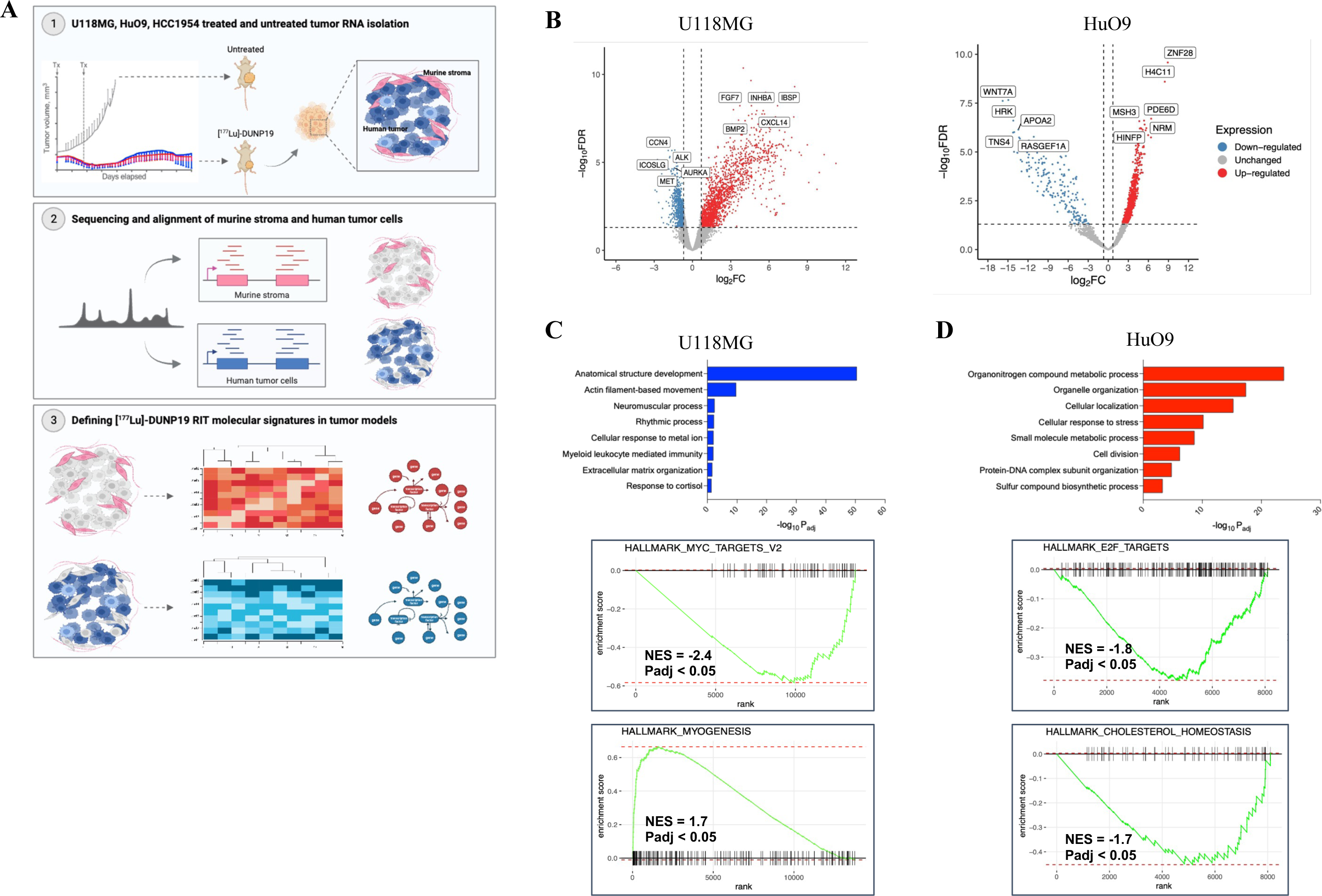
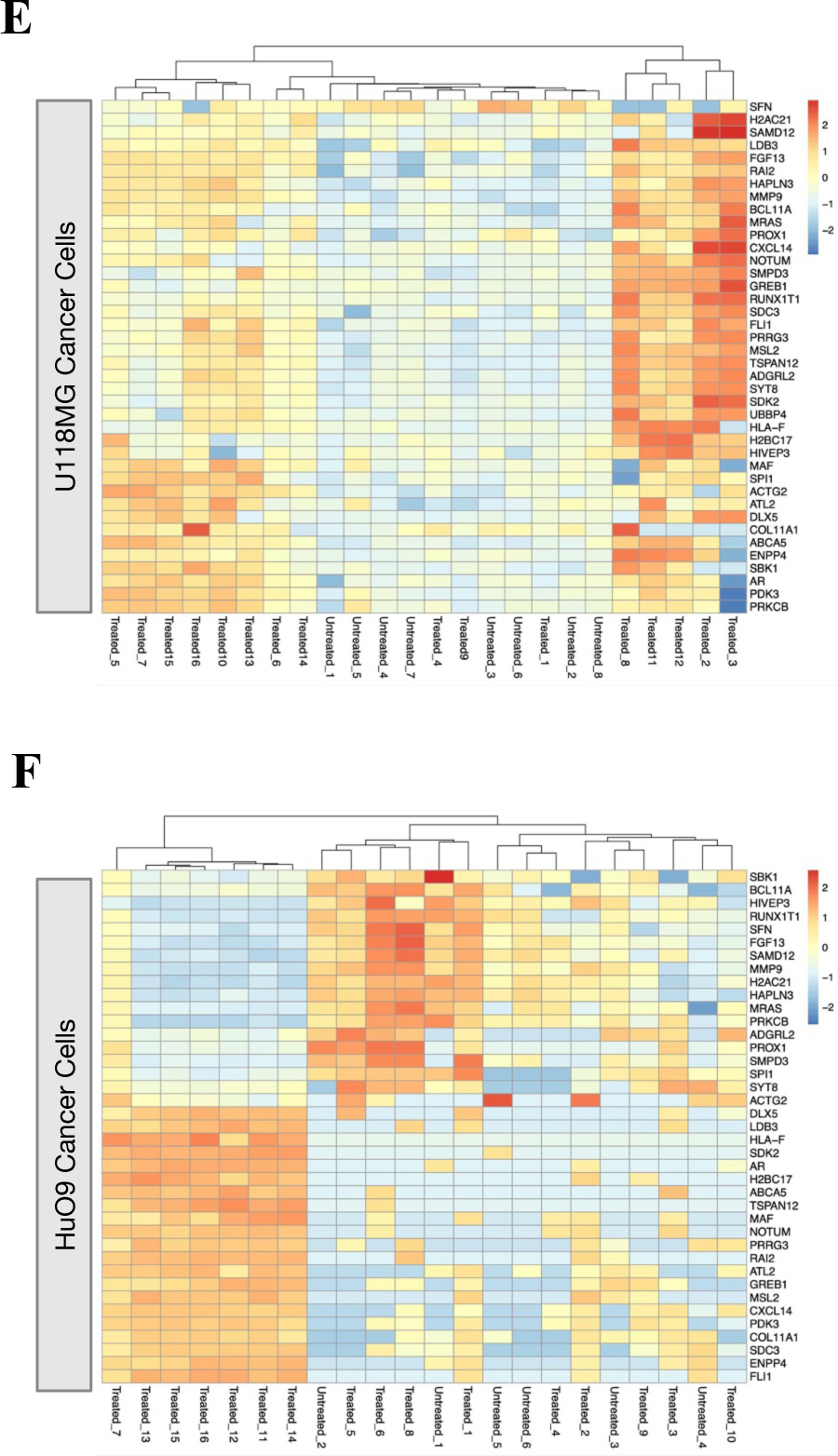
[^177^Lu]-DUNP19 induced radioimmunotherapy signatures in LRRC15+ cancer cells. **A.** Schematic of transcriptomic analysis of HuO9, U118MG, and HCC1954 tumors after [^177^Lu]-DUNP19 treatment. Treated or untreated tumor samples were harvested for RNA isolation, before sequencing and alignment to either murine or human genomes. Overlapping or ambiguous reads were discarded. **B.** Volcano plot of the top up-(red) and downregulated (blue) DEGs after treatment with [^177^Lu]-DUNP19 in U118MG (left) and HuO9 (right) cancer cells (FDR < 0.05). DEGs were ranked by fold-change. The top and bottom genes were labeled. **C, D.** Gene ontology (GO) biological pathway enrichment analysis of DEGs in treated **(C)** U118MG and **(D)** HuO9 cancer cells (adjusted p-value < 0.05). Enriched biological pathways with more than 10 overlapping terms (genes) were plotted by Padj value to indicate processes most significantly enriched after [^177^Lu]-DUNP19 RIT. **E, F.** Overlapping differentially expressed genes (40 genes, FDR < 0.05, FC > 1) in **(E)** U118MG and **(F)** HuO9 cancer cells after [^177^Lu]-DUNP19 treatment were plotted for visualization of [^177^Lu]-DUNP19-induced changes. Relative expression per gene was plotted to indicate up-(in red) or downregulated (blue) genes by Z-score normalization.

Due to the fractionated dosing regimens utilized in our therapeutic studies (Fig. 3, 4), our first objective was to determine whether there were significant differences in tumor gene signatures when comparing [^177^Lu]-DUNP19 dosing regimens. However, we did not find gene signatures that were associated with the [^177^Lu]-DUNP19 radiation dose received. Therefore, tumor samples were not separated by [^177^Lu]-DUNP19 radiation dose received in subsequent analyses and for each model, [^177^Lu]-DUNP19-treated tumors were compared to untreated tumors.

In human cancer cells, RNA-sequencing analysis identified 19,578 protein-coding genes, of which 1,985 (HuO9), 1,043 (U118MG), and 23 (HCC1954) were differentially expressed genes (DEGs) after [^177^Lu]-DUNP19 therapy. In [^177^Lu]-DUNP19-treated murine stroma, 663 (HuO9), 319 (U118MG), and 73 (HCC1954) DEGs were identified compared to untreated controls (Fig. 5B). In U118MG tumors, the most upregulated genes included bone morphogenic protein (*BMP2*) and inhibin subunit beta A (*INHBA*), two modulators of TGFβ signaling and epithelial-mesenchymal transition (Fig. 5B). In HuO9 tumors, epigenetic modulators such as *HINFP* and *H4C11* were upregulated, while anti-apoptotic genes (*HRK*) and genes in the canonical pro-tumorigenic WNT and RAS pathways were among the most significantly downregulated genes after DUNP19 RIT (Fig. 5B).

To gain a sense of the biological pathways altered by DUNP19 RIT, gene ontology analysis of DEGs from tumors containing LRRC15+ cancer cells (U118MG, HuO9) was performed (Fig. 5C,D). In treated U118MG tumors, gene ontology terms were immune- and epithelial-mesenchymal transition related (response to cortisol, myeloid leukocyte mediated immunity, anatomical structural development) (Fig. 5C). Further analysis using gene set enrichment analysis (GSEA) of genes ranked by fold-change (log_2_FC) revealed enrichment of myogenesis (myoblast stem cell differentiation) and a downregulation of MYC target genes (Fig. 5C, Supp. Table 1). In contrast, GSEA and gene ontology analysis of treated HuO9 tumors revealed alterations in metabolic and cell cycle-related pathways, as indicated by negative enrichment of E2F cell cycle transcription factors, as well as alterations in nitrate metabolism, lipoprotein homeostasis, and response to stress (Fig. 5F, Supp. Table 1). Despite differences in the biological pathways involved in DUNP19 RIT response, 40 genes were differentially expressed in both U118MG and HuO9 treated tumors (Fig. 5E,F). This was surprising given the different origins and different cell lineages of the two models. Of the 40 shared genes, several had functions similar to the hypothesized roles of LRRC15 (cell migration, invasion, and adhesion) or had been shown to be co-expressed with *LRRC15*, including *COL11A1*, *FGF13*, and *CXCL14* (Fig. 5E,F) (8, 21).

To complement our understanding of transcriptomic responses by DUNP19 RIT in the tumor microenvironment, we also conducted a comprehensive analysis of the murine stroma in HuO9, U118MG, and HCC1954 tumors. In all three models, RIT induced changes in pathways related to immune activation, including the upregulation of *Gzmk*, *Cxcr6*, and *Lck* (Fig. 6A-C). Whole transcriptome principal component analysis identified three distinct transcriptional clusters for treated cancer cells in the HuO9 and U118MG models and two clusters in HCC1954 (Supp. Fig. 7). Based on these results, we studied whether these changes could be explained by an overall shift in cell types present within the [^177^Lu]-DUNP19 treated tumor samples (i.e. loss of mesenchymal phenotypes). We employed Syllogist to further assess the proportional distribution of cell types in treated tumors compared to untreated samples (22). In accordance with the expression of *LRRC15* in cancer cells originating from mesenchymal stem cells, HuO9 and U118MG tumors displayed a notable overrepresentation of mesenchymal cells across all examined samples, with a significant loss of the mesenchymal cell phenotype in DUNP19-treated HuO9 cancer cells (Supp. Fig 8). No discernible alterations in relative cell composition were observed following treatment with [^177^Lu]-DUNP19 in other tumor models. Additionally, our observations revealed that HCC1954 tumor cells predominantly maintained an epithelial phenotype, consistent with the absence of *LRRC15* expression in the cancer cells (Supp. Fig 8).

**Figure 6.**
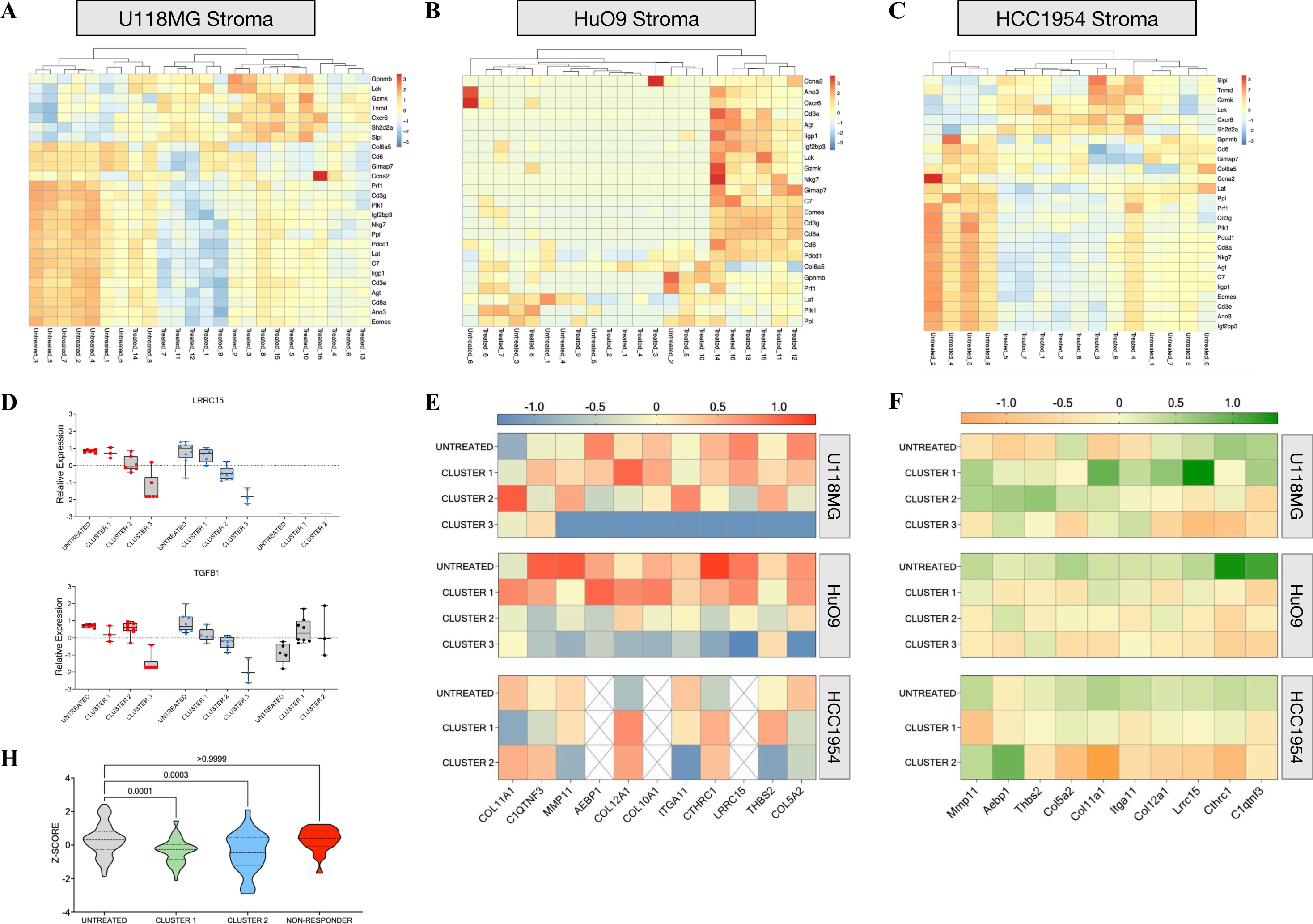
[^177^Lu]-DUNP19 therapy eradicates a LRRC15+ TGFβ signature associated with immunotherapy resistance A-C. DEGs overlapped in [^177^Lu]-DUNP19-treated stroma from U118MG (**A**, 26 genes), HuO9 (**B**, 23 genes), and HCC1954 (**C**, 26 genes) tumors. Relative expression (Z-score normalization) was plotted to indicate upregulated (red) or downregulated (blue) genes. **D.** Box- and-whisker plots representing relative transcript expression of *LRRC15* (top) and *TGFB1* (bottom), comparing untreated tumors to tumors after [^177^Lu]-DUNP19 therapy. HuO9 transcripts are plotted in red (left), U118MG in blue (middle), and HCC1954 in black (right). Samples were separated by transcript signature based on PCA plots and hierarchical clustering (Supp. Fig. 7) into 2 (HCC1954) or 3 (U118MG, HuO9) clusters. Expression of *LRRC15* and *TGFB1* in treated samples from cluster 3 are significantly (p<0.005) decreased in U118MG and HuO9, while no changes are observed in the LRRC15-HCC1954 cancer cells. **E, F.** Transcript data from clustered (Supp. Fig. 7) cancer cells **(E)** or tumor stroma **(F)** show decreased expression of the LRRC15+ TGFβ signature. **E.** Untreated U118MG (top) and HuO9 (middle) cancer cells lose expression of the LRRC15+ TGFβ signature after [^177^Lu]-DUNP19 treatment (red = high, blue = low expression). **F.** Loss of the LRRC15+ TGFβ signature is observed across all tumor stroma after [^177^Lu]-DUNP19 RIT (green = high, orange = low expression) **G.** HCC1954 tumors that were resistant to [^177^Lu]-DUNP19 treatment (defined as reaching 1000m^3^ endpoint before conclusion of study) had no significant reduction of the 11-gene LRRC15+ TGFβ signature within tumor stroma.

Overall, and in line with plasma membrane associated LRRC15 protein levels (Fig. 1B,C), *LRRC15* expression was higher in HuO9 than in U118MG cancer cells, and not quantifiable in HCC1954 cancer cells. In contrast, stromal *Lrrc15* expression was 3- and 4-fold higher in HCC1954 tumors than in U118MG and HUO9 tumors, respectively (Supp. Fig. 9). Comparison of *LRRC15/Lrrc15* expression in cancer cells and stroma of HuO9 and U118MG tumors across clusters showed a trend of decreased expression with increasing cluster distance from untreated samples in both cancer cells and stroma (Fig. 6D); the expression of *TGFB1*, a known regulator of LRRC15 expression, mirrored the *LRRC15/Lrrc15* expression pattern (Fig. 6D). Interestingly, *Lrrc15* and *Tgfb1* levels in HCC1954 stroma remained constant, while *TGFB1* expression was increased in treated LRRC15-null HCC1954 cancer cells.

Further analysis of LRRC15+ clusters showed that treatment with [^177^Lu]-DUNP19 resulted in the progressive loss of a gene signature associated with immune cell exclusion and poor response to immune checkpoint blockade in TGFβ-driven, LRRC15+ CAFs (4, 8) (Fig. 6E,F). Consistently, in two mice with HCC1954 tumors which progressed despite [^177^Lu]-DUNP19 RIT, neither *Lrrc15* nor the LRRC15+ CAF gene-signature decreased (Fig. 6H). Taken together, these findings collectively imply that LRRC15 serves as a critical modulator of the immune system within the tumor microenvironment and suggest that targeting LRRC15 with [^177^Lu]-DUNP19 could remodel the tumor microenvironment, hindering tumor progression.

## DISCUSSION

LRRC15 has emerged as a promising TGFβ-driven biomarker expressed on the cell membrane of cancer cells derived from mesenchymal stem cells and on a subset of cancer-associated fibroblasts within the tumor microenvironment (4, 8). Studies evaluating genes associated with metastatic progression have characterized LRRC15’s role in metastases to bone in breast cancer and to bowel in ovarian cancer, while LRRC15 knockdown by siRNA significantly inhibits tumor progression in preclinical models (23). Furthermore, a retrospective study assessing primary osteosarcoma lesions identified a correlation between LRRC15 expression, aggressive disease, and shorter overall survival (9).

The novel technology presented here harnesses the unique characteristics of a humanized IgG_1_ antibody, DUNP19, to specifically bind to a phylogenetically stable epitope of LRRC15. The rapid cellular internalization of DUNP19 exhibited by LRRC15-expressing cells provides an optimal foundation for leveraging DUNP19 as an effective vehicle for delivery of radionuclides to target cells. In this investigation, we have also substantiated DUNP19’s versatility by demonstrating its functionalization with both diagnostic and therapeutic radioisotopes. The radiolabeled modality facilitates identification of patients harboring LRRC15+ lesions, allowing for precise therapeutic intervention via molecularly specific PET or SPECT imaging. Subsequently, patients with LRRC15+ tissues can be selected for personalized therapeutic dosing, delivering tumor-specific ionizing radiation with minimal off-target effects. A similar approach is currently used in clinical practice, with examples including the application of radioactive iodine in thyroid cancer therapy and the use of radioligands binding to specific membrane antigens, such as [^177^Lu]-PSMA-617 for prostate-specific membrane antigen and [^177^Lu]-DOTATATE for somatostatin receptors. Patients exhibiting high tumoral uptake of the diagnostic radioligand on PET/CT imaging, reflecting elevated target expression and successful drug delivery, are deemed eligible for treatment with these therapeutic radioligands.

Our LRRC15-RIT approach exhibited effective targeting across various cancer models, including models of breast cancer, osteosarcoma, and glioblastoma, all representing highly lethal cancers with distinct tumor biology and LRRC15 expression patterns. Notably, the specific depletion of LRRC15+ cancer and stromal cells through a single systemic administration of [^177^Lu]-DUNP19 significantly slowed tumor progression and conferred a survival benefit in all models. These outcomes align with previous observations where the genetic ablation of LRRC15+ CAFs in murine models of pancreatic adenocarcinoma significantly reduced tumor volume and slowed tumor growth (8). Moreover, our findings underscore the practicality of directing therapeutic efforts towards LRRC15-deficient tumor cells via the crossfire phenomenon. Upon binding of [^177^Lu]-DUNP19 with adjacent LRRC15-expressing CAFs, the decay of ^177^Lu can instigate DNA breaks spanning multiple cell diameters. These crossfire effects can significantly improve responses in tumors with low or heterogeneous target expression (24, 25). The strategic application of a ^177^Lu antibody-conjugate addresses the challenge of heterogeneous LRRC15 expression within tumor tissue, thereby enhancing the therapeutic efficacy of the treatment. DUNP19 also binds to an epitope shared by human and murine LRRC15 and we do not expect the antibody to demonstrate vastly different biodistribution upon translation to patients. Moreover, *in vivo* cellular internalization, as observed for the DUNP19-based radioconjugate, has been reported to enhance radioisotope retention and reduce extracellular shedding of the radioisotope (26). In agreement with other beta-emitting RIT, we noted a decrease in white and red blood cells after administration of [^177^Lu]-DUNP19 (27). However, the bone marrow recovered within an expected time frame after treatment injection, allowing for serial dosing.

The LRRC15 protein is representative of a distinct set of TGFβ-driven genes that are predictors of immune checkpoint blockade resistance and unfavorable tumor evolution (4, 8). Given LRRC15’s known association with TGFβ, immunosuppression, and the observed potential immunomodulatory effects of RIT (28), our second aim was to explore whether the anti-tumor activity induced by [^177^Lu]-DUNP19 could reverse the signaling profile associated with immunosuppression and resistance to immunotherapy. To target tumor-supporting immunosuppressive stroma, many investigations have focused on fibroblast activation protein-α (FAP), a membrane bound serine protease overexpressed by CAFs (29). However, normal tissues and healthy activated fibroblasts also express FAP during wound healing, such as the activation of myofibroblasts post-myocardial infarction. While both LRRC15 and FAP have been shown to be upregulated by TGFβ (30), recent studies revealed a unique subset of TGFβ-driven CAFs that express LRRC15 and are negative for FAP expression (4, 8), and that these LRRC15+ CAFs are associated with CD8+ T-cell exclusion (8). During our therapeutic studies, a notable observation emerged: LRRC15+ tumors exhibited a concurrent reduction in both growth and the TGFβ–LRRC15 gene signature upon exposure to RIT. However, within a small subset (2 out of 12) of LRRC15-breast cancer (HCC1954) cases, tumor volumes did not decrease following the administration of [^177^Lu]-DUNP19. Moreover, this subset exhibited a persistent TGFβ–LRRC15 signature, suggesting a nuanced response within this specific context.

We also show that LRRC15-targeted RIT leads to differential expression of genes in stromal cells related to immune cell function, including genes indicative of activation and proliferation of T-cells (*Cd3, Cd8, Gimap7, Gzmk, Lat, Lck, Eomes*) and natural killer cells (*Cxcr6, Eomes, Prf1*). In the stroma of U118MG tumors, the T-cell suppressor gene encoding PD-1 was downregulated, suggesting that [^177^Lu]-DUNP19 may contribute to relief of immune cell suppression and T-cell exclusion, and that immune-based adjuvant therapies may be complementary to LRRC15-RIT. The transcriptomic remodeling of immune-related signaling pathways and expression of immune activating cytokines after RIT may increase immune cell activation and infiltration in previously immunologically “cold” tumors. Additionally, ablation of radio-resistant stroma may further increase immune cell invasion and help amplify the anti-tumor immune response, but future studies in immunocompetent mouse models will be required to investigate [^177^Lu]-DUNP19 RIT’s effects on the adaptive immune system and potential synergies with immunotherapies.

In the broader context of cancer treatment, our findings provide a novel technique for non-invasive imaging and treatment of a wide range of aggressive tumors with limited options for targeted therapy. Critically, we demonstrate that [^177^Lu]-DUNP19 therapy can be effective despite heterogenous LRRC15 expression within tumor tissue and that targeted radiation by [^177^Lu]-DUNP19 treatment is not limited to LRRC15+ tumor cells, taking advantage of the crossfire effects of beta-radioimmunotherapy. In addition, LRRC15’s regulation by TGFβ allows for successful targeting of pro-tumorigenic TGFβ signaling mechanisms that have been shown to contribute to immunotherapy resistance and poor prognosis. We show, at a transcriptomic level, that these TGFβ-LRRC15 signatures are largely erased in [^177^Lu]-DUNP19-treated tumors and expression of other anti-tumor immune pathways increases. Based on these observations, we hypothesize that the eradication of this TGFβ-LRRC15 signature in tumor cells would be synergistic with existing immunotherapies and allow for immune cell infiltration, as demonstrated by Krishnamurty et al (8). However, this remains an open question, as more advanced syngeneic models will be required to answer questions about immune functionality after LRRC15-targeted RIT. In summary, we propose targeting LRRC15+ cells with [^177^Lu]-DUNP19 as a novel theranostic strategy that provides sustained tumor control across models of LRRC15+ disease, improves survival, and reprograms the transcriptomic landscape of pro-tumorigenic and immunosuppressive mechanisms within the TME, all with minimal side effects.

## MATERIALS AND METHODS

### Reagents

All reagents were purchased from Sigma-Aldrich unless otherwise stated.

### DUNP19

For DUNP19 production, HEK293 cells were cultured in a 2L suspension using FreeStyle 293 Expression Medium (Life Technologies, Carlsbad, CA, USA) with a cell density maintained at 1 × 10^6 cells/mL on the day of transfection. Expression plasmids harboring genes for DUNP19 heavy chain and light chain in human IgG1/kappa format were combined with the transfection agent and incubated for 10 min at room temperature. The transfected cell culture was then incubated at 37 °C, 8% CO2 on an orbital shaker rotating at approximately 110 rpm for seven days. Culture medium was collected, subjected to centrifugation, and filtered through 0.22 µm filter systems. Antibodies were isolated through Protein A chromatography, followed by buffer exchange to PBS (pH 7.4) via gel filtration. The concentration was calculated from absorbance measurement at 280 nm.

For labeling with Lutetium-177 and Copper-64, respectively, DUNP19 was functionalized with benzylisothiocyanate derivatives of the acyclic chelating agent CHX-A’’-DTPA (p-SCN-CHX-A’’-DTPA) or cyclic NOTA (p-SCN-NOTA), respectively, using amine-reactive chemistry. Radiolabeling was performed as previously described (see Supplementary Materials and Methods) (31). The average radiochemical yield was 95±5 % with a radiochemical purity >99 % and an average specific activity of 4 MBq/mg. Injection solutions of [^177^Lu]-DUNP19 were formulated in 2% BSA/PBS containing 0.14 mM EDTA (100-fold molar excess). Binding affinity of the radio-conjugate to life cells was determined using Ligandtracer technology (32).

An amine-based protein labeling kit (Invitrogen, #A20173) was used for labeling DUNP19 with AlexaFluor-647 or AlexaFluor-594. Labeling was performed as described by the kit’s recommended protocol.

### Cell Culture

U118MG (glioblastoma), U87MG (glioblastoma), RPMI7951 (melanoma), NCI-H196 (small cell lung cancer), HCC1954 (breast cancer), SAOS2 (osteosarcoma), U2OS (OS), Kasumi-2 (leukemia), Calu-1 (non-small-cell lung cancer), RCH-ACV (leukemia), MHH-Call-3 (leukemia), Hs737.T (giant cell sarcoma), HEK293T, LNCaP (prostate cancer), and K7M2 (murine osteosarcoma) were purchased from ATCC. HuO9 (osteosarcoma) was purchased from the Japanese Cancer Research Resources Bank (Tokyo, Japan). All cell lines were cultured according to the manufacturer’s instructions and frequently tested for Mycoplasma.

To overexpress LRRC15, HEK293T and K7M2 cells were transduced with a pLenti-LRRC15-GFP-Puro vector (Origene, NM_130830) with a multiplicity of infection of 5. Optimal puromycin concentrations for stable selection of pLenti-LRRC15-GFP-Puro were determined by kill curves. Following selection with puromycin for 14 days, single cell clones were plated by serial dilution, expanded, and sorted for the top 1% of GFP+ cells.

### In Vitro Studies

#### Pulldown Assay (Immunoprecipitation -Mass Spectrometry)

Pierce Protein G Magnetic Beads (ThermoFisher, #88847) were pre-incubated with 70 ug DUNP19 antibody or hIgG1 mAb in PBS at room temperature (RT) for 1 h. Bead-antibody conjugate was recovered using magnetic separation before adding 300 ug protein lysates (isolated from U118MG cells) for 2 h on ice. Beads were washed with PBS and an on-bead digestion was performed with 0.25 % Trypsin for 16 h. MS-MS was run using the Agilent 6530 LC/MS in collaboration with the UCLA Molecular Instrumentation Center according to previously described protocols (33, 34).

#### Flow Cytometry

To assess the binding affinity of DUNP19, cells were blocked (10% normal goat serum in PBS, 15 minutes at RT) and incubated with serial dilutions of fluorophore-conjugated DUNP19 or hIgG1 mAb (0.051 – 1000 ng/mL) in triplicate for 1 h at RT. Cells were washed with 1% BSA in PBS (180 x g, 3 minutes) before adding viability dye per manufacturer’s instructions (Invitrogen, #L34989). Antibody binding capacity of cells was assessed using 1 µg/mL DUNP19 and anti-human IgG Simply Cellular bead standards (Bangs Laboratories, #816). Quantity of LRRC15 surface antigens available for DUNP19 binding was normalized to cell surface area (determined experimentally by confocal microscopy). All flow cytometry experiments were run in collaboration with UCLA’s Jonsson Comprehensive Cancer Center Flow Cytometry Shared Resource using an Attune NxT Flow Cytometer (Invitrogen). Flow cytometry data were analyzed using FlowJo (Version 10, BD Biosciences).

#### Reverse Transcriptase Polymerase Chain Reaction (RT-PCR)

Expression of *LRRC15* in cells was determined by Taqman qRT-PCR. Cells were lysed and one-step reverse transcription/quantitative PCR performed with Cells-to-CT Taqman kit (A25603). *LRRC15* was probed with Applied Biosystems Taqman Assay probes (Assay ID Hs00370056_s1) and normalized using *GAPDH* housekeeping gene (Assay ID Hs02786624_g1). All qPCR assays were run using the ViiA 7 Real-Time PCR system (Applied Biosystems).

#### Confocal Microscopy of Cells

For immunocytochemistry, cells were blocked with 10% normal goat serum in PBS for 1 h at room temperature. Cells were stained with DUNP19 (20 ug/mL) and anti-PMCA1 antibody (Abcam, ab3528, 1:100). Primary antibodies were incubated overnight at 4 °C before washing and staining with goat anti-rabbit AlexaFluor647 (1:500, Invitrogen A21235) and goat anti-human AlexaFluor488 (1:500, Invitrogen A11013) secondary antibodies. Cells were fixed with 3.7% paraformaldehyde for 15 minutes at RT before washing and mounting cells with Vectashield mounting media with DAPI (H1200-10) onto slides for imaging with a Leica TCS SP8 Digital Microscope.

To confirm internalization, cells (0.0015 x 10^6^ cells/well) were seeded in phenol red-free complete media in 384-well u-clear flat bottom black plates (Greiner, #781092) and stained with 1 ug/mL DUNP19-AF647, anti-huLAMP1-A488 antibody (1:250, Invitrogen, 53-1079-42) and Hoechst 33342 (1:2000 dilution, Invitrogen) at 4°C for 1 h to prevent antibody internalization while promoting surface binding. Unbound antibody was aspirated and replaced with phenol red-free complete media. Time-lapse confocal imaging was performed using a temperature controlled ImageXpress MicroXL High Content Imaging microscope (Molecular Devices). The microscope temperature was set to mimic cell culture conditions (37°C, 5% CO2) and images were taken at 10x magnification (4 sites per well) every 20 minutes for 6-12 h. The plasma membrane signal of DUNP19-AF647 relative to the cytosolic signal (LAMP1) at time 0 (directly after incubation at 4°C) was quantified to calculate the fraction of internalized antibodies over time.

### *In Vivo* Studies

#### Animal Studies

All animal experiments were conducted in compliance with national legislation on laboratory animal protection and permitted by the local ethics committees for animal research at Washington University, Lund University, University of California, Los Angeles and University of Duisburg-Essen.

#### Subcutaneous Tumor Models

Athymic nude mice (BALB/cAnNRj-Foxn1 nu/nu; 6–8 weeks old, 20–25 g; Janvier) were inoculated with U118MG (5.8 x 10^6^ cells), SAOS2 (6 x 10^6^ cells), HuO9 (6 x 10^6^ cells), HCC1954 (4.9 x 10^6^ cells) or LNCaP (5 x 10^6^ cells) cells in a 200 μL (1:1 v/v) mixture of media with Matrigel via subcutaneous injection in the right flank. Tumors developed after 3 to 6 weeks. Tumor volume was estimated with caliper measurements twice weekly (V (mm^3^) = 0.5 x length x width^2^).

#### Orthotopic Osteosarcoma Model

Nineteen athymic nude mice (BALB/cAnNRj-Foxn1 nu/nu; males, 6-8 weeks old, 24.2±1 g; Janvier) were anesthetized, and the tibia of the right hind limb was punctured. HuO9 cells (1.5 x 10^6^ cells in 10 μL media) were injected into the cavity using a microvolume syringe with a 27-gauge needle. Bone wax (Surgical Specialties Corporation, #903) was applied to seal the punctured area and prevent exodus of implanted cells before the area was washed with saline. Tumor development was confirmed using ultra-high-resolution CT and [^177^Lu]-DUNP19 SPECT/CT imaging (nanoScan; Mediso, Budapest, Hungary).

#### Small-Animal Imaging

PET: Male mice (B6;129-Rag2tm1FwaII2rgtm1Rsky/DwlHsd) (n_=_5) bearing s.c. SAOS2 xenografts were intravenously (i.v.) injected with [^64^Cu]-DUNP19 (8.75 MBq, 100 µg DUNP19 in 100 μL 10 mM ammonium acetate). Dynamic PET images were acquired during the first hour post-injection (p.i.), followed by static scans (20 minutes each) at 12 h, 24 h, and 36 h using a microPET R4 rodent scanner (Siemens). To confirm specificity of the DUNP19 PET-signal, imaging with the bone-seeking PET-probe Fluorine 18-sodium fluoride ([^18^F]-NaF; 10 MBq i.v.) was performed in mice bearing SAOS2 tumors. SPECT: Mice with intratibial HuO9 xenografts were i.v. injected with [^177^Lu]-DUNP19 (20 MBq, 30 μg). Mice were scanned for 50-60 minutes under anesthesia (2-3% isoflurane) using a SPECT/CT device (nanoScan Mediso, Budapest, Hungary). Details on PET, SPECT and CT acquisition are provided in the Supplementary Materials and Methods.

#### [^177^Lu]-DUNP19 Biodistribution and Kinetics

To investigate the biodistribution of [^177^Lu]-DUNP19, mice with s.c. SAOS2, HuO9, U118MG, HCC1954, or K7M2 tumors, respectively, received 0.5 MBq [^177^Lu]-DUNP19 (SAOS2, U118MG, K7M2: 16 µg; HuO9, HCC1954: 30 µg) by i.v. injection and were sacrificed at predetermined time points (SAOS2: 6, 24, 48, 72, 168 and 336 hours p.i.; HuO9: 6, 24, 48, 72, 168 and 336 h p.i.; U118MG: 24, 48 and 72 hours p.i.; n=4 mice/time point, HCC1954 and K7M2: 72 h p.i.). To confirm targeting specificity, mice bearing both LRRC15+ U118MG (left flank) and LRRC15-LNCaP (right flank) xenografts were i.v. injected with either 0.5 MBq [^177^Lu]-hIgG1 mAb or [^177^Lu]-DUNP19 (n=4 mice) and sacrificed 48 hours later. To study the effect of the antibody mass amount on the biodistribution and tumor uptake, mice bearing HuO9 tumors (n=5 mice/group) were administered escalating DUNP19 doses (1, 10, 30, 100 and 300 μg/mouse at 0.5 MBq ^177^Lu; i.v.) and euthanized 72 h p.i.. In all studies, blood, tumors, and several normal tissues were collected, dried and weighed. The radioactivity contained in the respective tissue and reference standards (0.5 MBq [^177^Lu]-DUNP19) was quantified in a NaI(Tl) automated well counter (1480 WIZARD; Perkin Elmer). Decay-corrected data were expressed as percent injected activity per gram tissue (% IA/g).

#### Radiological Analysis of Tumor Calcification

Quantification of radiopacity was analyzed using ImageJ software (version 1.53). A brightness threshold of 41 was used to define the area of each tumor sample. The total area of the tumor was then quantified using ImageJ’s built-in area analysis program. Areas of calcification were defined by a brightness threshold of 81 and quantified in the same manner. Percent calcification was calculated by dividing the area of calcification over the area of the entire tumor.

#### Confocal Microscopy of Xenografts

Mice with HuO9, SAOS2 and HCC1954 tumors, respectively, were injected with 30 ug DUNP19-AF594 when tumor volume reached between 200-300 mm^3^. Sections were washed two times with PBS before permeabilization with 0.1 % Triton-X/ PBS and blocking for 1 h at RT with 10 % goat serum/ PBS. Sections were stained with an anti-LAMP1-A488 antibody (HuO9, SAOS2: anti-hLAMP1, 1:250, Invitrogen, #53-1079-42; HC1954: anti-mLAMP1, 1:250, Invitrogen, #53-1071-82, to detect the murine stroma) overnight at 4°C. The next day, slides were stained with Phalloidin-AF647 for 1 h at RT (1:2000, Invitrogen, #A22283) and mounted with Vectashield Antifade mounting media with DAPI (Vector Laboratories, H-2000-2). Confocal microscopy was done by the UCLA Advanced Light Microscopy and Spectroscopy Laboratory (ALMS) with a Leica TCS-SP8 microscope at 63x magnification and sequential imaging for far-red, orange, green and blue emitting dyes. Z-stack images were taken and Lightning deconvolution and 3D reconstruction (Leica Microsystems) were performed during post-processing.

#### Therapy Studies

Average tumor volume at the start of treatment was 181 ± 20 mm^3^ and the average animal weight was 25.2 ± 0.7 g. Mice bearing s.c. HuO9 or U118MG xenografts with LRRC15 expression in both tumor cells as well as tumor stroma, and HCC1954 xenografts with LRRC15 expression only in the tumor stroma, were randomized into groups of 10-12 animals receiving either [^177^Lu]-DUNP19 (16-30 μg, i.v.) or no treatment. HCC1954 model: Mice were treated with a single injection of either 10 MBq or 20 MBq [^177^Lu]-DUNP19. HuO9 model: Animals received three fractionations [^177^Lu]-DUNP19 at days 0, 32 and 75 resulting in a cumulative administered activity of 50 MBq (group 1: 10+20+10 MBq; group 2: 20+10+20 MBq). To investigate the impact of pre-therapeutic tumor volume on the efficacy of [^177^Lu]-DUNP19 in the HuO9 model, mice (n=6-10 mice/group) were administered 30 MBq [^177^Lu]-DUNP19 (30 µg, i.v.) when tumors reached a volume of 203±64 mm^3^ (group 1) or 504±152 mm^3^ (group 2). U118MG model: Mice were treated with two fractions [^177^Lu]-DUNP19 at days 0 and 34 and a cumulative activity of 20 MBq (10+10 MBq) or 30 MBq (20+10 MBq). Treatment efficacy was assessed by measuring tumor growth and time to a humane endpoint. Mice survival was analyzed by using a log rank test in GraphPad Prism. P values <0.05 were considered significant statistically. Tumors from a subset of mice were harvested 90-120 days p.i. (except U118MG: day 155 p.i.) and processed for RNA-sequencing (see below).

To evaluate the efficacy of [^177^Lu]-DUNP19 in a clinically relevant orthotopic osteosarcoma model, mice with intratibial HuO9 tumors were randomized (23 days post-tumor engraftment) to receive 20 MBq [^177^Lu]-DUNP19 (30 µg, i.v.; n=12) or PBS (n=7). Four days after treatment, [^177^Lu]-DUNP19 tumor uptake and presence of viable HuO9 tumor was assessed by SPECT/CT imaging. At day 163 post-first injection, mice received an additional 20 MBq [^177^Lu]-DUNP19 and were re-scanned to detect residual viable HuO9 tumor tissue. Mice were followed up for 190 days after the first [^177^Lu]-DUNP19 injection.

#### Toxicity of [^177^Lu]-DUNP19

Body weights and hematological toxicity and recovery were monitored in mice treated with [^177^Lu]-DUNP19. Blood samples were taken before and weekly after injection of [^177^Lu]-DUNP19 for 4 weeks p.i.. Samples (20 µL) were collected from the tail vein of awake, immobilized mice by piercing the vein with a needle (27G) and collecting blood in a K2EDTA-coated plastic micropipette. Blood cell counts were obtained using an Exigo Veterinary Hematology Analyzer (Boule Medical, Stockholm, Sweden).

#### Immunohistochemistry

LRRC15 expression was analyzed on formalin-fixed, paraffin embedded sections using the Dako REAL Peroxidase Detection System (Dako) according to the manufacturer’s instructions. Antigen retrieval was performed by heat-induced epitope retrieval using Tris/EDTA (pH 8.1). Sections were incubated with anti-hLRRC15 antibody [EPR8188(2)] (1:100; #ab150376, Abcam) for 1 h at room temperature.

#### Gene Expression Analysis

For RNA-sequencing, tumor tissues were harvested at 90-120 days (HuO9, HCC1954) or 155 days (U118MG) post-treatment, apart from untreated mice (harvested when tumor volume measured greater than 1000 mm^3^, in accordance with established endpoint protocols). Tumor tissue was preserved in RNA*later* stabilization solution (Invitrogen, AM7020) before RNA isolation with the Qiagen RNeasy kit (#74004). RNA quality control was assayed via TapeStation (Agilent) and stranded mRNA library preparation performed in accordance with Illumina protocols. Samples were sequenced on Illumina’s Novaseq platform to generate 50bp paired-end reads. Library preparation and sequencing was done with the help of UCLA’s Technology Center for Genomics and Bioinformatics (TCGB).

Raw read count RNA-sequencing data were generated from untreated and [^177^Lu]-DUNP19 treated HuO9 (untreated n=6, treated n=16), U118MG (untreated n=8, treated n=16) and HCC1954 (untreated n=8, treated n=10) tumors. Paired-end reads were aligned to either human (Hg38) or murine (Mm19) genome using the STAR method, as previously described (35). Ambiguous reads were discarded and FastQC analysis was utilized to confirm sequence quality. Low read count filtering was used to remove transcriptomic features for which fewer than 4 samples had at least 5 read counts of a gene, as described by the EdgeR differential analysis user guide (36). For each tumor model, principal component analysis based on log2 counts in RStudio Version 2023.06.1 was plotted. K-Clustering and heat maps were generated on log2-transformed read counts to visualize gene signatures in treated versus untreated samples. Differential expression analysis to identify differentially expressed genes was performed using EdgeR (Bioconducter, Version 3.40.2) using quasi-likelihood F-tests within the EdgeR program. A false discovery rate of 0.05 (adjusted using Benjamini-Hochburg methodology) and absolute log2-fold change >1 were selected as the cutoff for DEGs within this analysis. A positive fold-change represented upregulation and a negative fold change represented downregulation of gene expression in treated tumors. For comparison and visualization of gene expression between clustered samples, z-scores were calculated per gene. Pathway analysis was performed using gene set enrichment analysis and molecular signatures defined using the human and murine Molecular Signature Database Hallmark pathways (37). Additional analysis was performed using Gene Ontology (GO) Biological Pathways. For cell classification and identification, Syllogist (22) was used to identify signatures present from 43 cell types within normalized gene expression matrices. For murine cell classification, murine orthologs were matched to Syllogist cell signatures using g:Profiler.

### Statistical Analyses

Statistical analyses were conducted using Graphpad Prism software (Version 9.5.1). Data are expressed as mean and standard deviation. Statistical comparisons were performed using one-way ANOVA with Tukey multi-comparison tests and unpaired Student’s t-tests. A P-value of less than 0.05 was considered statistically significant.

### Data Availability

RNA-sequencing data are available from the Gene Expression Omnibus NCBI database upon publication.

## SUPPLEMENTAL METHODS

### Conjugation and radiolabeling of DUNP19

All buffers were treated with Chelex 100 resin (sodium form, Merck KGaA, Darmstadt, Germany) to remove any metal ions and filtered through a 0.22 μm filter before use. Prior to conjugation, DUNP19 was buffer exchanged using an AMICON Ultra-0.5-centrifugal filter device with a MWCO 30 kDa (Millipore, Burlington, MA, USA). DUNP19 (500 μg in 500 μl 0.07 M sodium borate, pH 9.3) was mixed with the bifunctional chelator p-SCN-CHX-A’’-DTPA (Macrocyclics, Texas, USA) in a 6:1 molar ratio. The mixture was extensively vortexed and incubated overnight at 38°C. The reaction mixture was then centrifuged for 10 minutes at 14000 x g using a 30 kDa filter to remove the excess non-bound chelator. The concentrate containing the CHX-A’’-DTPA-DUNP19 conjugate was recovered, and the buffer was adjusted to 0.2 M ammonium acetate (pH 5.5). The conjugate was aliquoted and kept at −20 °C until labeling. For labeling with ^177^Lu, 30-50 μg of the conjugate was mixed with 15-20 MBq ^177^LuCl_3_ (Curium, Sweden) and incubated at 38°C with continuous vortexing for 30 min. Thereafter the radiolabeled conjugate was purified, and buffer exchanged using AMICON 30 kDa filter. Radiochemical yield and purity of the radioconjugate were determined using silica-impregnated ITLC strips (150–771 DARK GREEN Tec-Control Chromatography strips, Biodex Medical Systems) eluted with 0.2 M citric acid and measured using the Cyclone Storage Phosphor System (PerkinElmer, Waltham, MA, USA). To reduce radiolysis, the final product was diluted with 2% BSA/PBS buffer (pH 7.4) and 100X molar excess of EDTA was added to scavenge any free metal ions.

To determine the maximal attainable specific activity for labeling DUNP19 with ^177^Lu, decreasing amounts of CHX-A’’-DTPA-DUNP19 (10, 5, and 2.5 g) were incubated at 38°C for 30 minutes with a fixed quantity of ^177^LuCl3 (10 MBq). The radiochemical yield was determined using SG-ITLC, as previously described.

Conjugation of p-SCN-Bn-NOTA chelator to DUNP19 was performed in a manner similar to that used in the CHX-A’’-DTPA conjugation described above. Copper-64 was produced at the Washington University in St. Louis School of Medicine Cyclotron facility. ^64^CuCl2 (20 mCi; 25 uL) was diluted with a 10-fold excess 0.1 M ammonium acetate (NH4OAc), pH 5.5 and then added to NOTA-conjugated anti-LRRC15 antibody. After mixing for 30 min at room temperature, the antibody conjugate was purified by gel chromatography (PD10) into 0.1 M HEPES buffer in saline. Purity was assessed by radioITLC (Bioscan AR2000 using samples spotted on Whatman paper in a running buffer of 50 mM DTPA (pH 5.5). Quantitative labeling with >99% radiochemical purity was observed.

## AUTHOR CONTRIBUTIONS

Claire M Storey: writing-original draft, writing-review and editing, data collection, formal analysis

Mohamed Altai: writing-original draft, writing-review and editing, data collection, formal analysis, conceptualization

Katharina Lückerath: writing-original draft, writing-review and editing, data collection

Wahed Zedan: data collection

Henan Zhu: formal analysis

Marija Trajkovic-Arsic: data collection, formal analysis

Julie Park: writing-review and editing, formal analysis

Norbert Peekhaus: writing-review and editing

Jens Siveke: writing-review and editing

Henrik Lilljebjörn: writing-review and editing

Diane Abou: data collection

Haley Marks: data collection

Enna Ulmert: data collection

Hans Lilja: writing-review and editing

Alexander Ridley: writing-review and editing

Marcella Safi: data collection

Constance Yuen: data collection

Susanne Geres: data collection

Liqun Mao: data collection

Michael Cheng: formal analysis

Johannes Czernin: writing-review and editing

Ken Herrmann: writing-review and editing

Laurent Bentolila: data collection, formal analysis

Xia Yang: writing-review and editing, formal analysis

Thoas Fioretos: writing-review and editing, formal analysis

Thomas Graeber: formal analysis

Kjell Sjöström: writing-review and editing

Robert Damoiseaux: writing-review and editing

Daniel Thorek: investigation, data collection, writing-review and editing

David Ulmert: writing-original draft, writing-review and editing, conceptualization

## ACKNOWLEDGEMENTS

This study was supported in part by the UCLA Eli and Edythe Broad Center of Regenerative Medicine and Stem Cell Research Rose Hill Foundation Innovator Award. The study was further supported by NCI R01CA201035, R01CA240711, R01CA229893, DoD W81XWH-18-1-0223, UCLA SPORE in Prostate Cancer (P50 CA092131), JCCC Cancer support grant from NIH P30 CA016042 (PI: Teitell), Knut and Alice Wallenberg Foundation, Bertha Kamprad Foundation, David H. Koch Prostate Cancer Foundation Young Investigator Award, Swedish Research Council, Swedish Cancer Society, SIPEA Foundation, Swedish Childhood Cancer Foundation, John and Augusta Perssons Foundation, Royal Physiographic Society of Lund, Franke and Margareta Bergqvist Foundation, Crafoord Foundation, Lund University Medical Faculty research time allocation award, and IngaBritt and Arne Lundberg Research Foundation. Confocal laser scanning microscopy was performed at the Advanced Light Microscopy/Spectroscopy Laboratory (RRID: SCR_022789) and the Leica Microsystems Center of Excellence at the California NanoSystems Institute at UCLA with funding support from NIH Shared Instrumentation Grant S10OD025017. Flow cytometry was performed in the UCLA Jonsson Comprehensive Cancer Center (JCCC) and Center for AIDS Research Flow Cytometry Core Facility that is supported by National Institutes of Health awards P30 CA016042 and 5P30 AI028697, and by the JCCC, the UCLA AIDS Institute, the David Geffen School of Medicine at UCLA, the UCLA Chancellor’s Office, and the UCLA Vice Chancellor’s Office of Research. We also thank the UCLA Technology Center for Genomics and Bioinformatics (TCGB) for preparation of the RNA sequencing. We express our gratitude to Joanna and Sven-Erik Strand for their assistance with intravenous tail-vein injections and administration, respectively, at Lund University. Additionally, we acknowledge the Lund University BioImaging Center (LBIC) for providing experimental imaging resources. We also acknowledge Hadis Westin and Jos Buijs from Ridgeview AB Sweden, for technical support with the LigandTracer technology. HL was supported in part from NIH/NCI (P30-CA008748, U01-CA266535, R01-CA244948) and Swedish Cancer Society (Cancerfonden 23 3074 Pj 01 H).

---

**Supplemental Figure 1.** DUNP19 binds to human and murine LRRC15 protein. **A.** DUNP19 binding kinetics to human (left) and murine (right) LRRC15 recombinant protein using the Bio-Layer Interferometry Octet system. Association of 100ug/mL DUNP19 (pink trace) or 100ug/mL IgG1 isotype control (green trace) are shown in real time over 850 s. Association and dissociation of DUNP19 to hLRRC15 or mLRRC15 is measured by binding rate (nm). **B.** The 11 unique proteins present after an immunoprecipitation-mass spectrometry (IP-MS) analysis of crude protein lysates from U118MG cells incubated with Protein G magnetic bead-conjugated DUNP19. Pulldown-MS composition was compared to protein composition present in non-specific IgG1 with U118MG lysates. LRRC15 is among the top proteins in complex with DUNP19.

**Supplemental Figure 2.** Live confocal microscopy-based assays to determine internalization kinetics of DUNP19 after chelator conjugation and in various cell types. **A.** AF647-DUNP19 or AF647-DUNP19-DOTA conjugate internalize into U118MG cells at similar rates. To determine internalization rate, images were taken every hour across 12 h at 37C. Time to 50% internalization (T_1/2_) of AF647-DUNP19 (measured by ratio of cytosolic intensity of AF647-DUNP19 compared to membrane integrated intensity) was 5.16 ± 0.83 h. The AF647-DUNP19-DOTA conjugate internalized at a similar rate, with a T_1/2_ of 5.75 ± 0.93 h. **B.** Internalization of AF647-DUNP19 in fibroblast-derived Hs819.T cells compared to osteosarcoma SAOS2 cancer cells. To determine Internalization rate was determined by an 8 h microscopy-based assay with images were taken every 10m at 37C. Time to 50% internalization (T_1/2_) of AF647-DUNP19 in SAOS2 cells was 1.31 ± 0.14 h. T_1/2_ internalization of AF647-DUNP19 into Hs819.T cells was significantly slower with a T_1/2_ of 1.66 ± 0.09 h.

**Supplemental Figure 3.** Expanded tumor biodistribution of [^177^Lu]-DUNP19 in U118MG, SAOS2, HuO9, K7M2*^LRRC15+^*and HCC1954 xenograft models across 6 timepoints; 6h, 24h, 48h, 72h, 168h, and 336h. Tumor accumulation of [^177^Lu]-DUNP19 peaks between 48-72 h for all models and is represented as %IA/g (percent injected activity per gram tissue). At 72h, tumoral accumulation is as follows; U118MG: 14.31 ± 2.01 %IA/g, SAOS2: 23.07 ± 2.90 %IA/g, HuO9: 43.93 ± 7.89 %IA/g, K7M2*^LRRC15+^*: 13.60 ±1.49 %IA/g, HCC1954: 11.83 ± 2.50 %IA/g.

**Supplemental Figure 4.** IHC and IF analysis of *ex vivo* LRRC15+ tumor models. **A.** *Ex vivo* immunohistochemistry analysis of HuO9, K7M2*^LRRC15+^*, and K7M2 wildtype tumor sections harvested from untreated animals. Images are stained for LRRC15 and counterstained with hematoxylin, shown at 4x and 20x. **B.** Confocal microscopy of tumor tissues obtained from animals treated with fluorescently labeled DUNP19. Confocal images of s.c. HuO9 (LRRC15+ cancer cells / LRRC15+ CAF) tumors harvested at 72 h post-i.v. injection of AF594-DUNP19 (yellow). Tumor sections were co-stained for Actin (red), DNA (DAPI, blue) and LAMP1 (lysosomal marker, green). Images show that DUNP19 accumulates in the cellular cytoplasm and co-localized with LAMP1 indicating intracellular trafficking of the mAb to the lysosomal compartments (arrow) after binding to LRRC15.

**Supplemental Figure 5.** Animal weights across tumor models (HuO9, U118MG, HCC1954) corresponding to therapy studies in Figure 3 (HuO9, top), and Figure 4 (U118MG, middle, and HCC1954, bottom). Gray arrows denote administration of [^177^Lu]-DUNP19 or PBS. Overall, weight was stable throughout [^177^Lu]-DUNP19 therapy studies across tumor models.

**Supplemental Figure 6.** Toxicity studies examining blood cell counts during treatment course across tumor models (HuO9, U118MG, HCC1954). HuO9 (top), U118MG (middle) and HCC1954 (bottom) exhibited transient reductions in lymphocytes and monocytes after administration of [^177^Lu]-DUNP19, which recovered to baseline levels within 3 weeks, recapitulating toxicity profiles observed in clinical cases.

**Supplemental Figure 7.** Principal component analysis (PCA) plots of whole transcriptome cancer cell reads (left) and stromal cell reads (right). PC1 and PC2 show separation of treated samples in distinct clusters that were determined to correlate with LRRC15 transcript responses to [^177^Lu]-DUNP19 RIT. In HuO9 and HCC1954 tumor models, treated cancer cells and stromal cells from the same sample clustered in similar patterns. U118MG and HuO9 treated tumors were grouped into three distinct clusters, whereas HCC1954 treated tumors formed two clusters that were differentiated from untreated mice.

**Supplemental Figure 8.** Relative cell characterization of individual tumor samples using Syllogist. Overall, 43 cell types were analyzed and relative expression was averaged into seven summary cell types as shown (Germ/ESC, Neural/Glial, Mesenchymal, Endothelial, Epithelial, HSPC, Myeloid, Lymphoid). Normalized relative expression of cell types is presented on a scale of 0 (underrepresented cell type, not present in sample) to 1 (overrepresented cell type, present in sample). U118MG and HuO9 cancer cells were characterized as mesenchymal phenotypes, whereas HCC1954 cancer cells were majority epithelial (left plots), in line with relative LRRC15 cancer cell expression between the three models. In stromal cells, no significant cell phenotype was observed in any of the three models (right plots).

**Supplemental Figure 9.** Stromal cell expression of *Lrrc15* and *Tgfb1*. **A.** Transcript data from clustered (Supp. Fig.7) tumor stroma of *Lrrc15* (top) and *Tgfb1* (bottom) relative expression, represented by box- and-whisker plots. HuO9 transcripts are plotted in green (left), U118MG in orange (middle), and HCC1954 in grey (right). In HuO9 and U118MG stromal cells, expression of *Tgfb1* is significantly reduced in cluster 3 (p<0.05). Across tumor models, changes in *Lrrc15* transcript expression were not significant between treated and untreated samples. **B.** Comparison of relative baseline transcript expression of *Lrrc15* in tumor stroma, comparing HCC1954 (right), HuO9 (middle) and U118MG (left). In HCC1954 tumors, median stromal *Lrrc15* expression is high (8.91) compared to moderate expression in HuO9 (3.02) and U118MG (3.48).

**Table 1.** Biodistribution of [^177^Lu]-DUNP19 in male BALB/c-nu/nu mice bearing SAOS2 osteosarcoma xenografts 6-336 h after intravenous injection. The measured radioactivity of different organs is expressed as %ID/g and presented as an average value from 4 animals ± SD.

|  | **6 h** | **24 h** | **48 h** | **72 h** | **168 h** | **336 h** |
| --- | --- | --- | --- | --- | --- | --- |
| **Blood** | 18.84±1.12 | 11.86±1.67 | 9.31±0.66 | 8.74±1.08 | 5.07±1.02 | 2.58±0.63 |
| **Salivary gland** | 2.3±0.34 | 3.37±0.67 | 2.82±0.86 | 3.06±0.54 | 1.92±0.2 | 1.07±0.21 |
| **Heart** | 5.8±0.77 | 3.88±0.68 | 3.16±0.75 | 2.84±0.45 | 1.44±0.27 | 0.65±0.19 |
| **Lung** | 7.29±0.64 | 5.48±1.15 | 4.18±0.84 | 4.32±0.86 | 2.98±0.65 | 1.54±0.39 |
| **Liver** | 7.55±3.35 | 4.95±1.15 | 3.62±0.52 | 3.42±0.65 | 3.09±0.88 | 1.56±0.32 |
| **Spleen** | 4.99±1.05 | 4.41±0.68 | 4.54±1.19 | 3.76±0.7 | 2.79±0.77 | 2.02±0.28 |
| **Stomach** | 1.68±0.23 | 1.09±0.67 | 1.12±0.31 | 0.93±0.06 | 0.55±0.39 | 0.4±0.1 |
| **Small Intestine** | 2.2±0.38 | 1.58±0.32 | 1.42±0.51 | 1.31±0.25 | 0.74±0.25 | 0.37±0.12 |
| **Colon** | 1.94±0.54 | 1.13±0.2 | 0.75±0.14 | 0.91±0.2 | 0.5±0.08 | 0.27±0.09 |
| **Kidney** | 6.43±0.6 | 4.34±0.29 | 3.65±0.53 | 3.29±0.12 | 2.13±0.42 | 1.04±0.21 |
| **Tumor** | 3.86±0.86 | 18.09±1.41 | 17.87±3.04 | 23.1±2.9 | 20.55±6.19 | 15.64±4.1 |
| **Skin** | 3.07±1.06 | 6.07±1.1 | 6.2±0.73 | 7.64±1.23 | 4.65±1.63 | 2.5±0.48 |
| **Muscle** | 0.77±0.13 | 1.02±0.26 | 0.86±0.07 | 0.78±0.13 | 0.56±0.15 | 0.28±0.06 |
| **Bone** | 1.71±0.2 | 1.62±0.17 | 1.62±0.36 | 1.6±0.26 | 1.32±0.19 | 0.96±0.26 |
| **Brain** | 0.61±0.18 | 0.52±0.21 | 0.25±0.09 | 0.33±0.08 | 0.23±0.15 | 0.11±0.04 |

**Table 2.** Biodistribution of [^177^Lu]-DUNP19 in male BALB/c-nu/nu mice bearing HuO9 osteosarcoma xenografts 6-336 h after intravenous injection. The measured radioactivity of different organs is expressed as %ID/g and presented as an average value from 4 animals ± SD.

|  | **6 h** | **24 h** | **48 h** | **72 h** | **168 h** | **336 h** |
| --- | --- | --- | --- | --- | --- | --- |
| **Blood** | 23.81±3.1 | 14.69±2.7 | 13.68±1.1 | 11.56±1.1 | 4.49±1.4 | 1.74±0.52 |
| **Heart** | 6.6±1.3 | 4.31±1.14 | 4.36±0.97 | 3.9±0.34 | 1.51±0.45 | 0.67±0.19 |
| **Lung** | 10.58±0.9 | 7.69±1.21 | 6.6±0.67 | 5.89±0.67 | 2.62±0.41 | 1.43±0.33 |
| **Liver** | 9.83±2.58 | 6.52±0.81 | 6.15±0.17 | 5.78±0.58 | 2.95±0.12 | 2.62±0.96 |
| **Spleen** | 7.9±1.37 | 10.35±1.7 | 9.69±1.24 | 8.18±1.68 | 4.09±2.35 | 3.36±1.59 |
| **Stomach** | 2.34±0.31 | 1.75±0.41 | 1.53±0.25 | 1.43±0.1 | 0.7±0.33 | 0.31±0.15 |
| **Small Intestine** | 2.84±1.12 | 2.02±0.35 | 1.86±0.05 | 1.9±0.34 | 0.82±0.54 | 0.4±0.13 |
| **Colon** | 2.77±0.49 | 1.9±0.39 | 2.66±2 | 1.75±0.38 | 0.59±0.18 | 0.28±0.08 |
| **Kidney** | 9.25±1.25 | 6.68±1.32 | 4.65±2.21 | 5.35±0.84 | 2.48±0.27 | 0.9±0.13 |
| **Tumor** | 9.46±3.52 | 24.75±4.7 | 34.1±11.7 | 43.93±7.9 | 23.9±12.6 | 31.53±7.6 |
| **Skin** | 4.92±1.52 | 9.09±2.03 | 10.25±1.3 | 7.47±1.56 | 7.43±1.38 | 2.62±0.82 |
| **Muscle** | 1.39±0.13 | 1.52±0.35 | 1.39±0.28 | 1.26±0.2 | 0.83±0.2 | 0.45±0.1 |
| **Bone** | 2.35±0.38 | 2.57±0.39 | 2.36±0.09 | 2.63±0.7 | 1.38±0.51 | 0.91±0.23 |
| **Brain** | 0.78±0.34 | 0.4±0.09 | 0.44±0.11 | 0.29±0.09 | 0.13±0.03 | 0.06±0.02 |

**Table 3.** Biodistribution of [^177^Lu]-DUNP19 in male BALB/c-nu/nu mice bearing U118MG glioblastoma xenografts 24, 48 and 72 h after intravenous injection. The measured radioactivity of different organs is expressed as %ID/g and presented as an average value from 4 animals ± SD. *\*GI tract uptake is presented as %ID only*.

|  | **24 h** | **48 h** | **72 h** |
| --- | --- | --- | --- |
| **Blood** | 9.63±0.77 | 7.57±1.39 | 5.75±0.73 |
| **Salivary gland** | 1.74±0.35 | 1.92±0.14 | 1.89±0.21 |
| **Heart** | 2.40±0.50 | 2.10±0.55 | 1.74±0.14 |
| **Lung** | 4.60±0.77 | 4.21±0.88 | 3.34±0.64 |
| **Liver** | 6.22±3.00 | 4.79±0.74 | 6.17±1.19 |
| **Spleen** | 4.61±1.83 | 4.79±1.22 | 4.03±1.13 |
| **Stomach** | 0.71±0.27 | 0.56±0.29 | 0.71±0.15 |
| **Kidney** | 3.82±0.54 | 2.92±0.37 | 2.40±0.24 |
| **Tumor** | 12.04±0.92 | 14.31±2.01 | 13.30±1.08 |
| **Muscle** | 0.65±0.27 | 1.14±0.25 | 0.96±0.32 |
| **Bone** | 0.44±0.30 | 1.35±0.63 | 0.97±0.23 |
| **Brain** | 0.12±0.06 | 0.18±0.03 | 0.28±0.07 |
| **GI tract\*** | 2.63±0.34 | 2.24±0.26 | 2.33±0.31 |

**Table 4.** Biodistribution of [^177^Lu]-DUNP19 in male BALB/c-nu/nu mice bearing murine K7M2*^LRRC15+^* osteosarcoma and human HCC1954 breast cancer xenografts after intravenous injection. The measured radioactivity of different organs is expressed as %ID/g and presented as an average value from 4 animals ± SD. *\*GI tract uptake is presented as %ID only*.

| K7M2<sup>LRRC15+</sup> |  |  | HCC1954 |  |
| --- | --- | --- | --- | --- |
|  | 48 h | 72 h |  | 72 h |
| **Blood** | 7.62±0.76 | 6.98±1.13 | **Blood** | 6.95±1.67 |
| **Lung** | 3.80±0.37 | 4.55±1.19 | **Lung** | 4.01±1.03 |
| **Liver** | 4.67±1.21 | 4.88±1.22 | **Liver** | 4.37±0.72 |
| **Spleen** | 3.15±0.84 | 3.04±0.82 | **Spleen** | 3.97±0.49 |
| **Stomach** | 0.81±0.11 | 0.91±0.27 | **Stomach** | 0.81±0.08 |
| **Small Intestine** | 0.99±0.04 | 1.05±0.18 | **Small Intestine** | N/A |
| **Colon** | 0.72±0.14 | 0.98±0.24 | **Colon** | N/A |
| **Kidney** | 2.90±0.30 | 2.76±0.32 | **Kidney** | 2.77±0.35 |
| **Tumor** | 10.46±0.69 | 13.60±1.49 | **Tumor** | 11.83±2.5 |
| **Skin** | 4.88±0.52 | 6.43±1.45 | **Skin** | 6.04±1.06 |
| **Muscle** | 0.83±0.12 | 1.12±0.59 | **Muscle** | 0.95±0.36 |
| **Bone** | 1.21±0.28 | 1.17±0.40 | **Bone** | 1.16±0.12 |
| **Brain** | N/A | 0.26±0.07 | **Brain** | 2.29±0.31 |

## REFERENCES

1. Jin S, Sun Y, Liang X, Gu X, Ning J, et al. Emerging new therapeutic antibody derivatives for cancer treatment. Signal Transduct Target Ther 2022;7:39

2. Bejarano L, Jordão MJC, Joyce JA. Therapeutic Targeting of the Tumor Microenvironment. Cancer Discov 2021;11:933–959

3. Van der Heide CD, Dalm SU. Radionuclide imaging and therapy directed towards the tumor microenvironment: a multi-cancer approach for personalized medicine. EJNMMI 2022; 49:4616–4641

4. Dominguez CX, Muller S, Keerthivasan S, Koeppen H, Hung J, Gierke S, et al. Single-Cell RNA Sequencing Reveals Stromal Evolution into LRRC15(+) Myofibroblasts as a Determinant of Patient Response to Cancer Immunotherapy. Cancer Discov 2020;10:232–53

5. Purcell JW, Tanlimco SG, Hickson J, Fox M, Sho M, Durkin L, et al. LRRC15 Is a Novel Mesenchymal Protein and Stromal Target for Antibody-Drug Conjugates. Cancer Res 2018;78:4059–72

6. Wang Y, Liu Y, Zhang M, Lv L, Zhang X, Zhang P, Zhou Y. LRRC15 promotes osteogenic differentiation of mesenchymal stem cells by modulating p65 cytoplasmic/nuclear translocation. Stem Cell Res Ther 2018;9:65

7. Yang Y, Wu H, Fan S, Bi Y, Hao M, Shang J. Cancer_associated fibroblast_derived LRRC15 promotes the migration and invasion of triple_negative breast cancer cells via Wnt/beta_catenin signalling pathway regulation. Mol Med Rep 2022;25

8. Krishnamurty AT, Shyer JA, Thai M, Gandham V, Buechler MB, Yang YA, et al. LRRC15(+) myofibroblasts dictate the stromal setpoint to suppress tumour immunity. Nature 2022;611:148–54

9. Cui J, Dean D, Wei R, Hornicek FJ, Ulmert D, Duan Z. Expression and clinical implications of leucine-rich repeat containing 15 (LRRC15) in osteosarcoma. J Orthop Res 2020;38:2362–72

10. Ghandi, M., Huang, F.W., Jané-Valbuena, J. et al. Next-generation characterization of the Cancer Cell Line Encyclopedia. Nature 2019;569, 503–08

11. McDevitt MR, Thorek DLJ, Hashimoto T, Gondo T, Veach DR, Sharma SK, et al. Feed-forward alpha particle radiotherapy ablates androgen receptor-addicted prostate cancer. Nat Commun 2018;9:1629

12. Kurth J, Krause BJ, Schwarzenbock SM, Stegger L, Schafers M, Rahbar K. External radiation exposure, excretion, and effective half-life in (177)Lu-PSMA-targeted therapies. EJNMMI Res 2018;8:32

13. Deng R, Meng YG, Hoyte K, Lutman J, Lu Y, Iyer S, et al. Subcutaneous bioavailability of therapeutic antibodies as a function of FcRn binding affinity in mice. MAbs 2012;4:101–9

14. Thorek DLJ, Watson PA, Lee SG, Ku AT, Bournazos S, et al. Internalization of secreted antigen-targeted antibodies by the neonatal Fc receptor for precision imaging of the androgen receptor axis. Sci Transl Med 2016;8:367

15. Shi Y, van der Meel R, Chen X, Lammers T. The EPR effect and beyond: Strategies to improve tumor targeting and cancer nanomedicine treatment efficacy. Theranostics 2020;10:7921–7924

16. Illidge TM, Bayne M, Brown NS, Chilton S, Cragg MS, Glennie MJ, et al. Phase 1/2 study of fractionated (131)I-rituximab in low-grade B-cell lymphoma: the effect of prior rituximab dosing and tumor burden on subsequent radioimmunotherapy. Blood 2009;113:1412–21

17. Schlom J, Molinolo A, Simpson JF, Siler K, Roselli M, Hinkle G, et al. Advantage of dose fractionation in monoclonal antibody-targeted radioimmunotherapy. J Natl Cancer Inst 1990;82:763–71

18. Linden O, Hindorf C, Cavallin-Stahl E, Wegener WA, Goldenberg DM, Horne H, et al. Dose-fractionated radioimmunotherapy in non-Hodgkin’s lymphoma using DOTA-conjugated, 90Y-radiolabeled, humanized anti-CD22 monoclonal antibody, epratuzumab. Clin Cancer Res 2005;11:5215–22

19. DeNardo GL, Schlom J, Buchsbaum DJ, Meredith RF, O’Donoghue JA, Sgouros G, et al. Rationales, evidence, and design considerations for fractionated radioimmunotherapy. Cancer 2002;94:1332–48

20. Kreel L, Bydder G. Tumor calcification after radiotherapy demonstrated by computed tomography. J Comput Tomogr 1980;4:245–9

21. Guderian RH, Chico ME, Rogers MD, Pattishall KM, Grogl M, Berman JD. Placebo controlled treatment of Ecuadorian cutaneous leishmaniasis. Am J Trop Med Hyg 1991;45:92–7

22. Lu IN, Dobersalske C, Rauschenbach L, Teuber-Hanselmann S, Steinbach A, et al. Tumor-associated hematopoietic stem and progenitor cells positively linked to glioblastoma progression. Nat Commun 2021;12(1):3895

23. Mariani A, Wang C, Oberg AL, Riska SM, Torres M, Kumka J, et al. Genes associated with bowel metastases in ovarian cancer. Gynecol Oncol 2019;154:495–504

24. Iravani A, Violet J, Azad A, Hofman MS. Lutetium-177 prostate-specific membrane antigen (PSMA) theranostics: practical nuances and intricacies. Prostate Cancer Prostatic Dis 2020;23:38–52

25. Seifert R, Kessel K, Schlack K, Weckesser M, Kersting D, et al. Total tumor volume reduction and low PSMA expression in patients receiving Lu-PSMA therapy. Theranostics 2021;11:8143–51

26. Edwards WB, Xu B, Akers W, Cheney PP, Liang K, Rogers BE, et al. Agonist-antagonist dilemma in molecular imaging: evaluation of a monomolecular multimodal imaging agent for the somatostatin receptor. Bioconjug Chem 2008;19:192–200

27. Bodei L, Kidd M, Paganelli G, Grana CM, Drozdov I, Cremonesi M, et al. Long-term tolerability of PRRT in 807 patients with neuroendocrine tumours: the value and limitations of clinical factors. Eur J Nucl Med Mol Imaging 2015;42:5–19

28. Kleinendorst SC, Oosterwijk E, Bussink J, Westdorp H, Konijnenberg MW, Heskamp S. Combining Targeted Radionuclide Therapy and Immune Checkpoint Inhibition for Cancer Treatment. Clin Cancer Res 2022;28:3652–7

29. Hofheinz RD, al-Batran SE, Hartmann F, Hartung G, Jager D, Renner C, et al. Stromal antigen targeting by a humanised monoclonal antibody: an early phase II trial of sibrotuzumab in patients with metastatic colorectal cancer. Onkologie 2003;26:44–8

30. Juillerat-Jeanneret L, Tafelmeyer P, Golshayan D. Regulation of Fibroblast Activation Protein-α Expression: Focus on Intracellular Protein Interactions. J Med Chem 2021;64(19):14028–14045

31. Westerlund K, Altai M, Mitran B, Konijnenberg M, Oroujeni M, Atterby C, et al. Radionuclide Therapy of HER2-Expressing Human Xenografts Using Affibody-Based Peptide Nucleic Acid-Mediated Pretargeting: In Vivo Proof of Principle. J Nucl Med 2018;59:1092–8

32. Bondza S, Foy E, Brooks J, Andersson K, Robinson J, Richalet P, Buijs J. Real-time Characterization of Antibody Binding to Receptors on Living Immune Cells. Front Immunol 2017;8:455

33. Have S, Boulon S, Ahmad Y, Lamond AI. Mass spectrometry-based immuno-precipitation proteomics -the user’s guide. Proteomics 2011;11:1153–9

34. Lagundzin D, Krieger KL, Law HC, Woods NT. An optimized co-immunoprecipitation protocol for the analysis of endogenous protein-protein interactions in cell lines using mass spectrometry. STAR Protoc 2022;3:101234

35. Dobin A, Davis CA, Schlesinger F, Drenkow J, Zaleski C et al. STAR: ultrafast universal RNA-seq aligner, Bioinformatics 2013;29:15–21

36. Robinson MD, McCarthy DJ, Smyth GK. edgeR: a Bioconductor package for differential expression analysis of digital gene expression data. Bioinformatics 2010;26:139–40

37. Liberzon A, Birger C, Thorvaldsdottir H, Ghandi M, Mesirov JP, Tamayo P. The Molecular Signatures Database (MSigDB) hallmark gene set collection. Cell Syst 2015;1:417–25

