## Supplemental Figures for "Development of a LRRC15-Targeted Radio-Immunotheranostic Approach to Deplete Pro-tumorigenic Mechanisms and Immunotherapy Resistance"

### Supplemental Figure S1

**A**

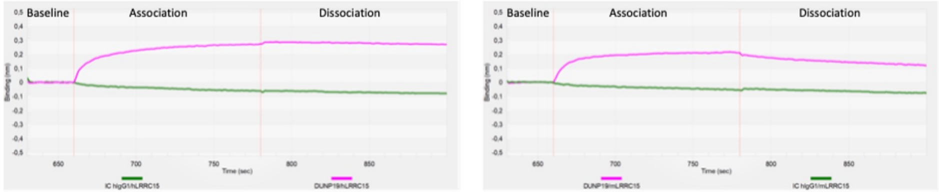


**B**

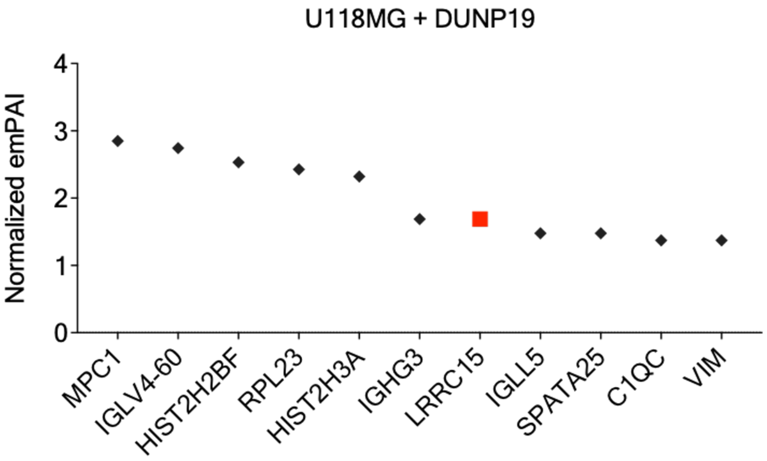

### Supplemental Figure S2

**A**

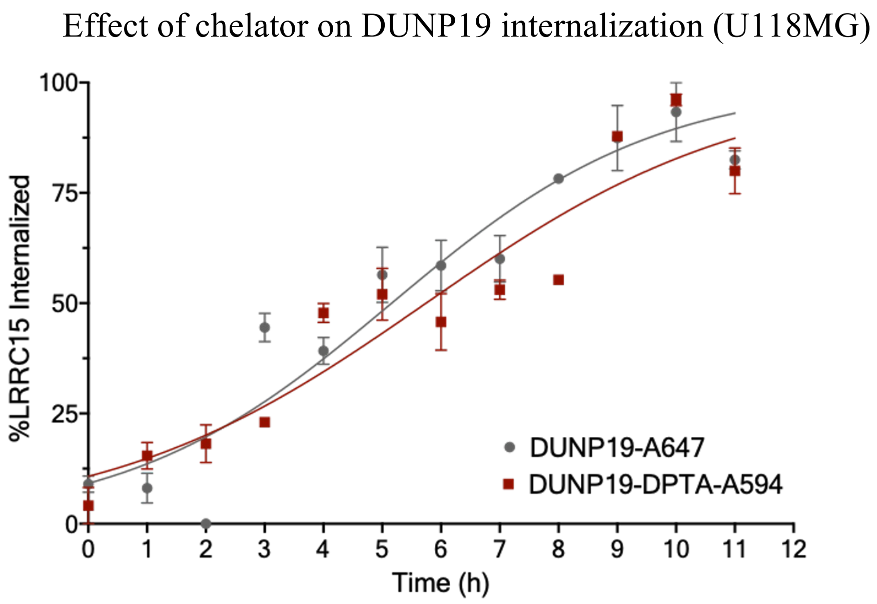


**B**

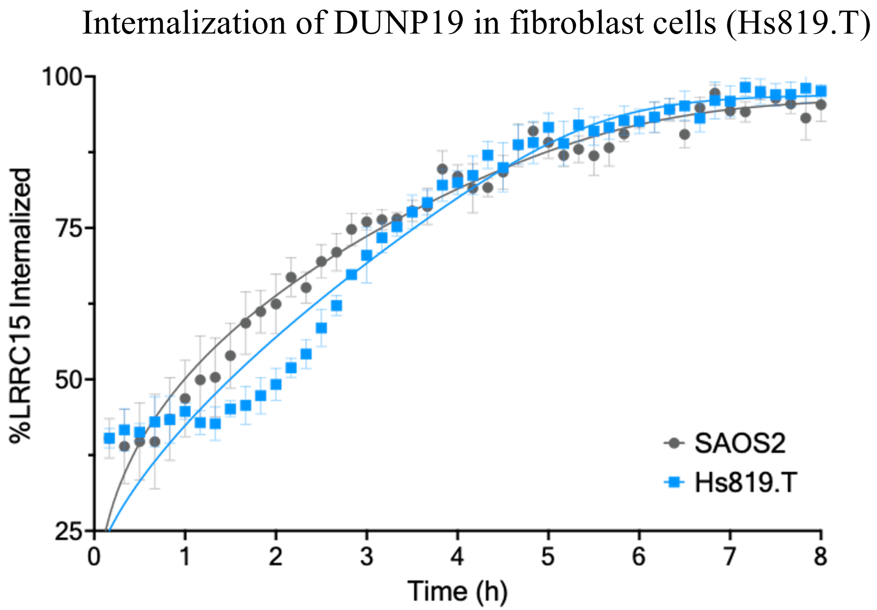

### Supplemental Figure S3

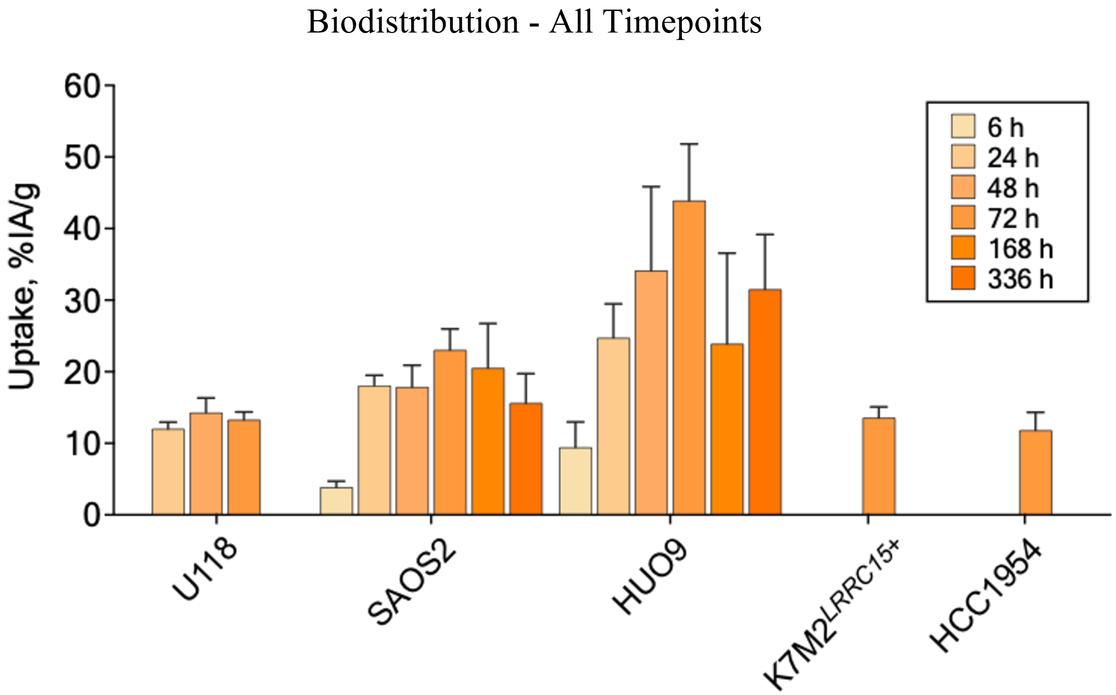

### Supplemental Figure S4

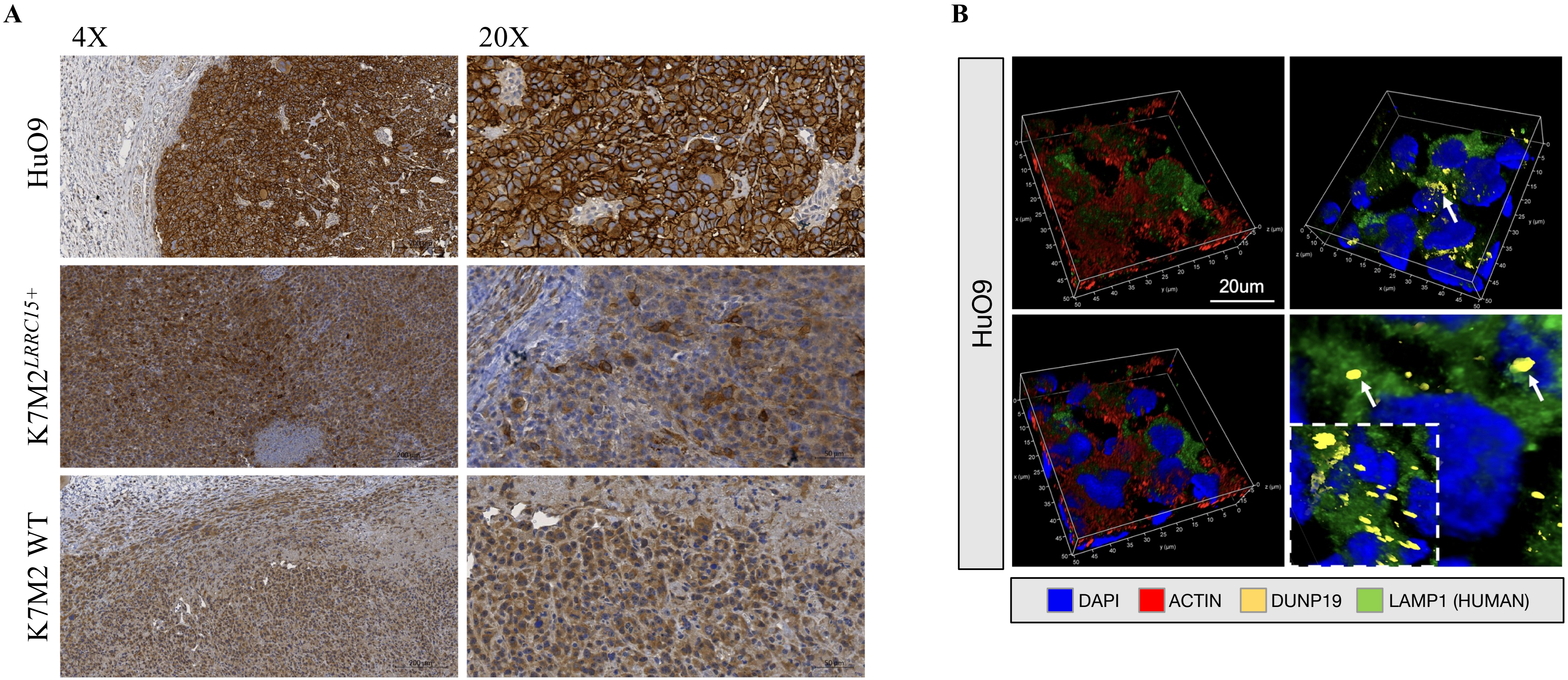

### Supplemental Figure S5

A

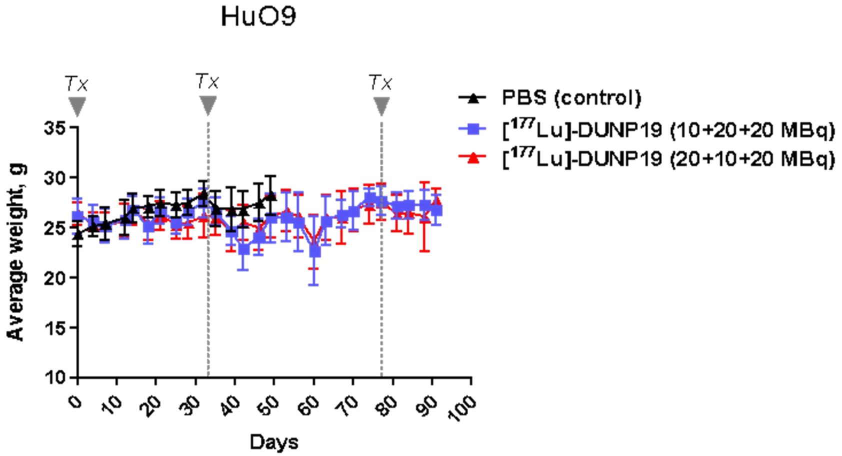


C

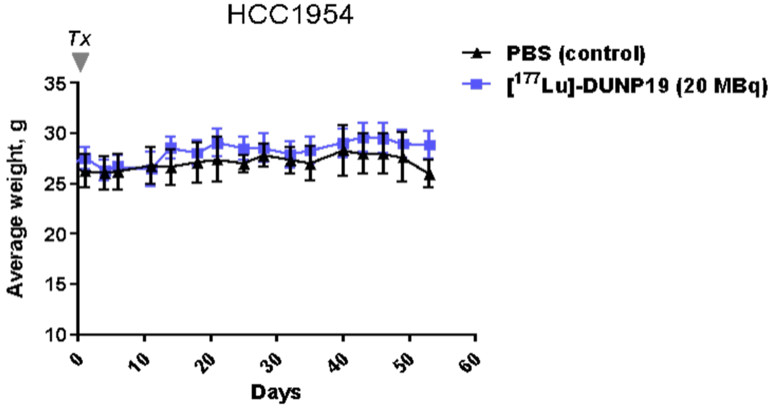


B

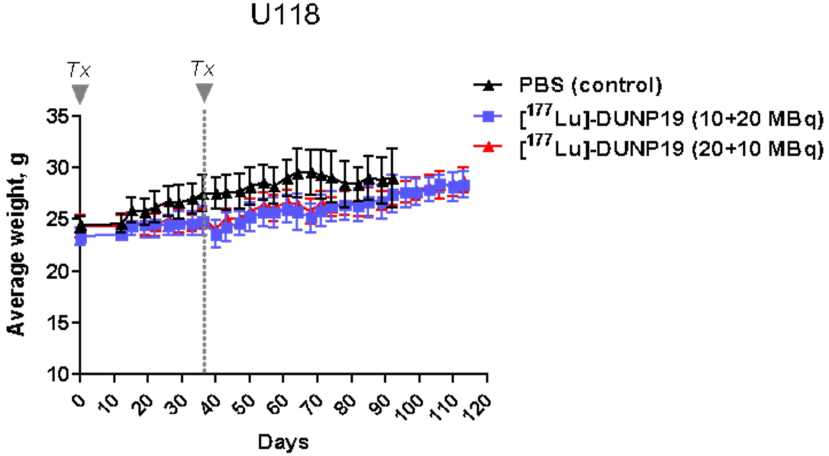

### Supplemental Figure S6

**A**

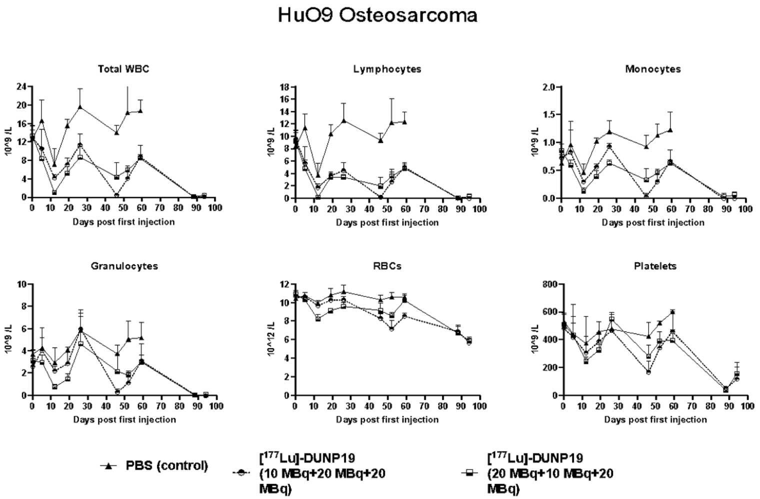


**B**

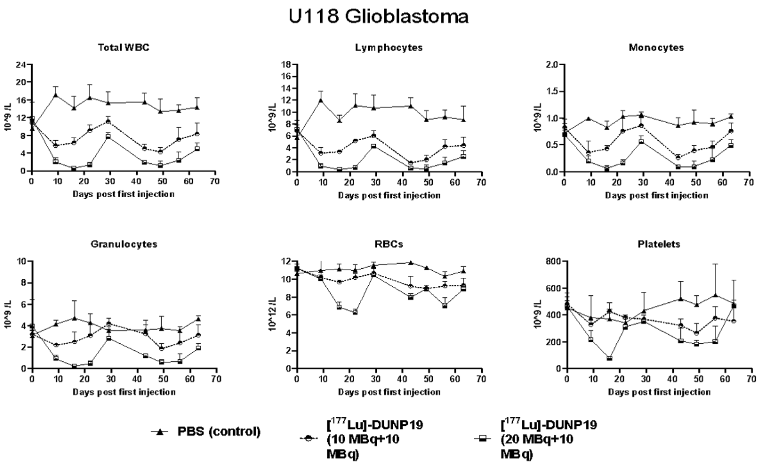


**C**

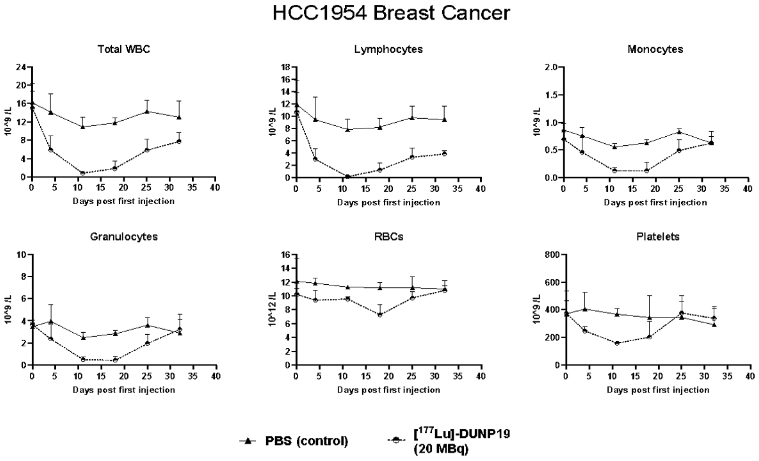

### Supplemental Figure S7

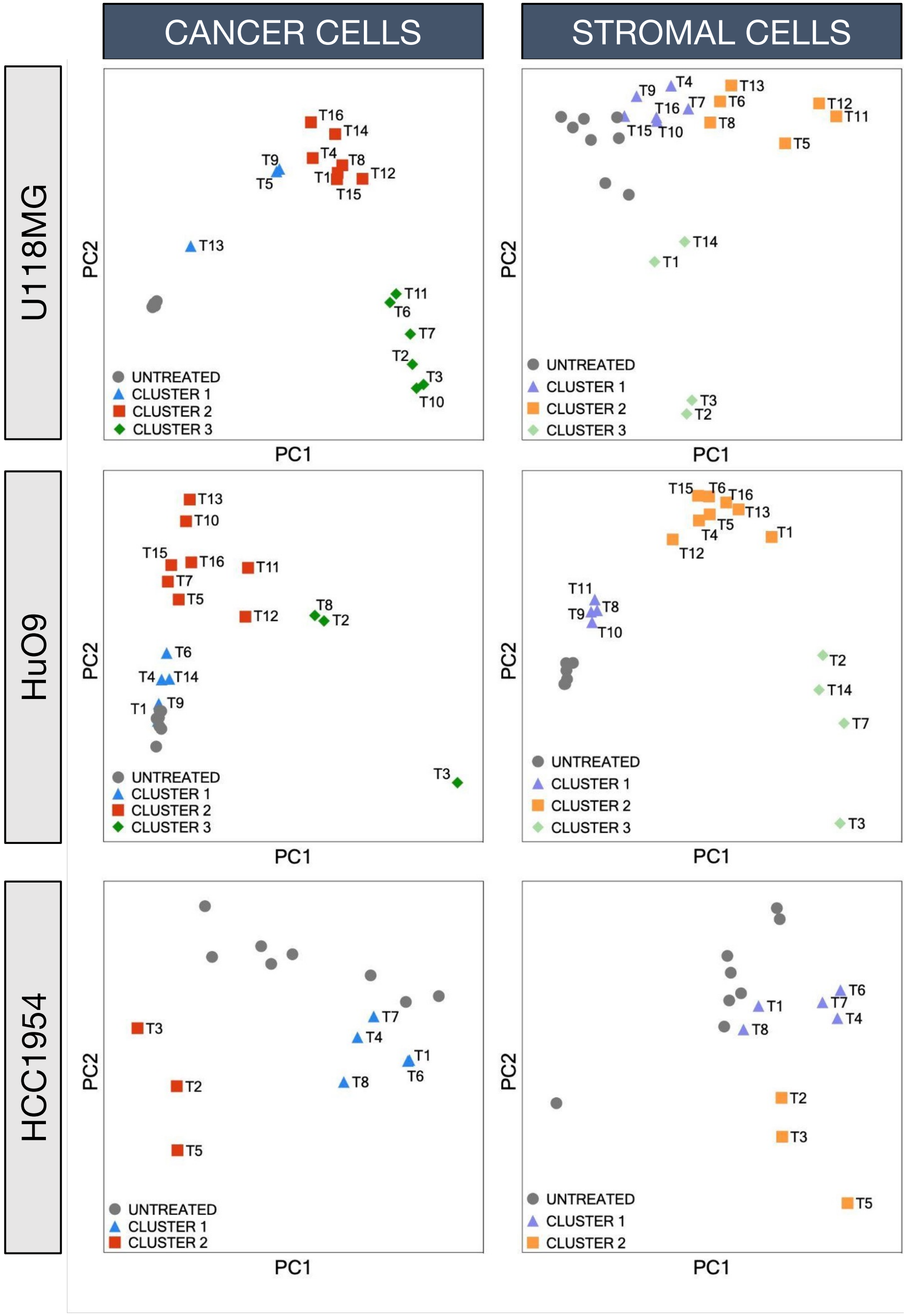

### Supplemental Figure S8

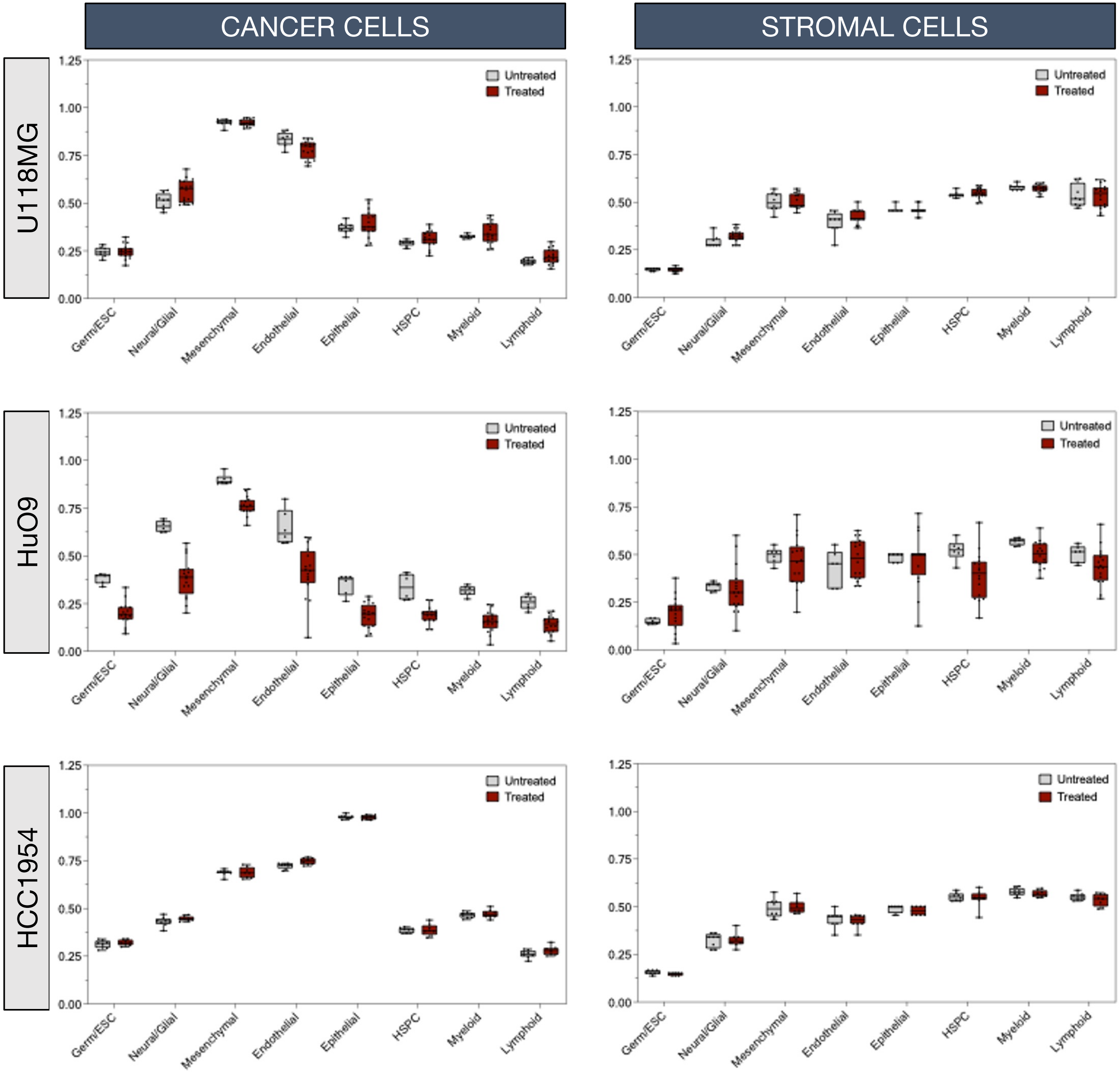

### Supplemental Figure S9

**A**

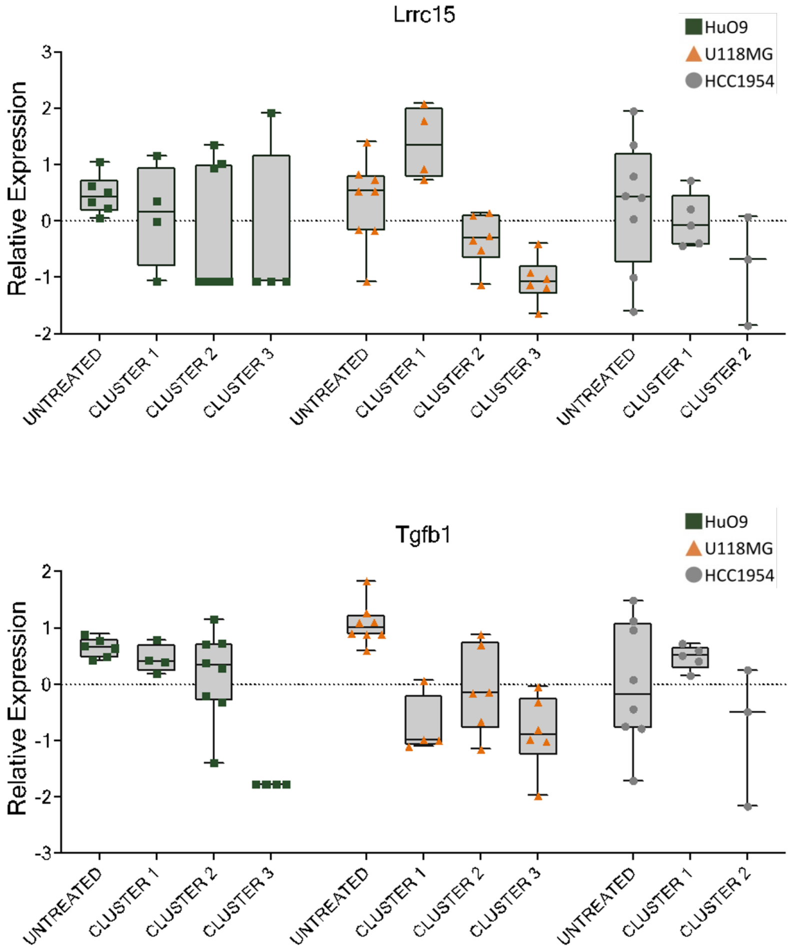


**B**

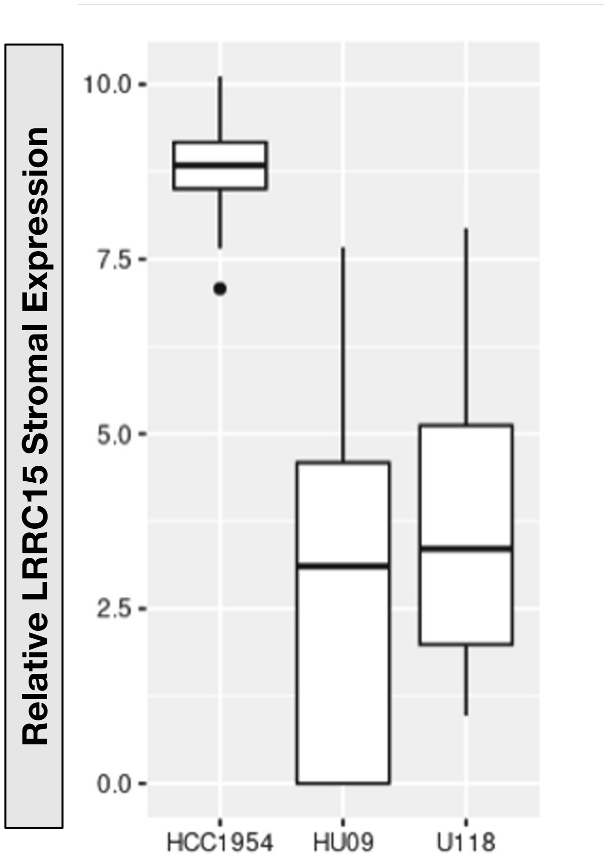

**Table S1.** Gene set enrichment analysis of HuO9, U118MG, and HCC1954 cancer cells

HuO9

| Pathway | Pval | Padj | Enrich Score | Norm Enrich Score |
|---|---|---|---|---|
| HALLMARK_E2F_TARGETS | 0.000 | 0.001 | -0.380 | -1.752 |
| HALLMARK_CHOLESTEROL_HOMEOSTASIS | 0.001 | 0.015 | -0.453 | -1.700 |
| HALLMARK_FATTY_ACID_METABOLISM | 0.002 | 0.015 | -0.392 | -1.643 |
| HALLMARK_OXIDATIVE_PHOSPHORYLATION | 0.001 | 0.015 | -0.351 | -1.595 |
| HALLMARK_GLYCOLYSIS | 0.001 | 0.015 | -0.362 | -1.584 |

U118MG

| Pathway | Pval | Padj | Enrich Score | Norm Enrich Score |
|---|---|---|---|---|
| HALLMARK_MYC_TARGETS_V2 | 0.000 | 0.000 | -0.583 | -2.406 |
| HALLMARK_MYOGENESIS | 0.000 | 0.000 | 0.664 | 1.745 |
| HALLMARK_TNFA_SIGNALING_VIA_NFKB | 0.000 | 0.000 | 0.623 | 1.637 |
| HALLMARK_KRAS_SIGNALING_DN | 0.000 | 0.000 | 0.666 | 1.698 |
| HALLMARK_KRAS_SIGNALING_UP | 0.000 | 0.000 | 0.613 | 1.607 |

HCC1954

| Pathway | Pval | Padj | Enrich Score | Norm Enrich Score |
|---|---|---|---|---|
| HALLMARK_INTERFERON_GAMMA_RESPONSE | 0.000 | 0.000 | -0.620 | -2.186 |
| HALLMARK_INTERFERON_ALPHA_RESPONSE | 0.000 | 0.000 | -0.695 | -2.286 |
| HALLMARK_OXIDATIVE_PHOSPHORYLATION | 0.000 | 0.000 | 0.446 | 2.063 |
| HALLMARK_GLYCOLYSIS | 0.000 | 0.000 | 0.425 | 1.945 |
| HALLMARK_INFLAMMATORY_RESPONSE | 0.000 | 0.000 | -0.554 | -1.903 |

**Table S2.** Gene set enrichment analysis of HuO9, U118MG, and HCC1954 stromal cells

| | Pathway | Pval | Padj | Enrich Score | Norm Enrich Score |
|---|---|---|---|---|---|
| HuO9 | HALLMARK_G2M_CHECKPOINT | 0.000 | 0.000 | -0.546 | -2.295 |
| | HALLMARK_E2F_TARGETS | 0.000 | 0.000 | -0.524 | -2.174 |
| | HALLMARK_INTERFERON_ALPHA_RESPONSE | 0.000 | 0.000 | -0.510 | -1.928 |
| | HALLMARK_MYOGENESIS | 0.000 | 0.000 | -0.428 | -1.782 |
| | HALLMARK_MYC_TARGETS_V1 | 0.000 | 0.001 | -0.404 | -1.724 |

| | Pathway | Pval | Padj | Enrich Score | Norm Enrich Score |
|---|---|---|---|---|---|
| U118MG | HALLMARK_E2F_TARGETS | 0.000 | 0.000 | -0.725 | -3.288 |
| | HALLMARK_G2M_CHECKPOINT | 0.000 | 0.000 | -0.692 | -3.154 |
| | HALLMARK_ALLOGRAFT_REJECTION | 0.000 | 0.000 | -0.694 | -3.046 |
| | HALLMARK_INTERFERON_GAMMA_RESPONSE | 0.000 | 0.000 | -0.671 | -2.987 |
| | HALLMARK_ADIPOGENESIS | 0.000 | 0.000 | 0.688 | 2.371 |

| | Pathway | Pval | Padj | Enrich Score | Norm Enrich Score |
|---|---|---|---|---|---|
| HCC1954 | HALLMARK_MYOGENESIS | 0.000 | 0.000 | -0.688 | -2.479 |
| | HALLMARK_G2M_CHECKPOINT | 0.000 | 0.000 | -0.669 | -2.424 |
| | HALLMARK_E2F_TARGETS | 0.000 | 0.000 | -0.640 | -2.317 |
| | HALLMARK_ALLOGRAFT_REJECTION | 0.000 | 0.000 | -0.614 | -2.182 |
| | HALLMARK_COAGULATION | 0.000 | 0.000 | 0.559 | 1.953 |
